# Multiomics dissection of the CO_2_-dependent fast growth of *Picochlorum celeri*

**DOI:** 10.64898/2026.09.27.753040

**Authors:** Yasin Torres-Tiji, Thayne Ekness, Devin Karns, Camilo Posso, Lye Meng Markillie, Kent Bloodsworth, Nathalie Munoz Munoz, Marina Gritsenko, Daniel MacLaughlin, Kaelyn Carpenter, Morgan K. Hampton, Natalie Furrer, Alaina LaPanse, Hugh Mitchell, Vladislav A. Petyuk, Niaz Bahar Chowdhury, Abby Jerger, Doo Nam Kim, James M. Fulcher, Rosalie Chu, Ronald Moore, Vanessa L. Paurus, Priscila Lalli, Josie Eder, Ernesto S. Nakayasu, Paul Piehowski, Angela Cintolesi, Jeremy Zucker, Alex Beliaev, Matthew Posewitz

## Abstract

With a sub-3-µm cell and a 27-Mbp diploid genome, the small but mighty *Picochlorum celeri* doubles within three hours and exceeds 30 g m^−2^ d^−1^ outdoors with CO_2_ supplied. In climate-simulating photobioreactors, productivity declined from 39 g m^−2^ d^−1^ under 2.25% CO_2_ to near zero in air and resumed within hours of CO_2_ restoration with photosystem II photochemistry largely retained. We profiled the transcriptome, proteome, phosphoproteome, ubiquitinome, acetylome, metabolome and lipidome, including the first site-resolved green-algal ubiquitinome. In air, the Rubisco large subunit tripled to 7.4% of protein while its two small-subunit isoforms, 70% identical, exchanged near-reciprocally, a novel CO_2_-dependent switch that we propose retunes Rubisco to low CO_2_. Photorespiratory enzymes increased in abundance whereas proteins involved in external nitrogen assimilation decreased, consistent with increased reliance on internal nitrogen recycling as carbon fixation became limiting. Transcript and protein responses were nearly uncoupled, with cytosolic ribosomal proteins declining despite increased transcripts and plastid ribosomal proteins spared. Phosphorylation was extensively remodeled, as were the lysine modifications ubiquitination, acetylation and CO_2_-dependent carbamylation, where a CO_2_-competition assay identified 86 bicarbonate-protected lysines, including RbcL K252. CO_2_ limitation therefore imposes a reversible growth arrest that reduces cytosolic translational investment while preserving photosynthetic capacity for rapid recovery.

## Introduction

Oxygenic photosynthesis is the engine that sustains most life on Earth, providing the energy and organic carbon that fuel ecosystems and meet humanity’s needs for food and biological resources. Microalgae contribute substantially to this global carbon flux, and their capacity for rapid growth makes them valuable systems for investigating the biological determinants of photosynthetic productivity (Torres-Tiji et al. 2020). Among microalgae, the marine chlorophyte *Picochlorum celeri* combines exceptionally fast growth with a small unicellular form and a remarkably compact diploid genome, with a haploid size of 13.7 Mbp—approximately an order of magnitude smaller than that of *Chlamydomonas reinhardtii* (Becker et al. 2020; Merchant et al. 2007). Comparative genomics indicates that genome streamlining in the *Picochlorum* lineage has involved the loss of specific metabolic pathways and reductions in gene copy number, suggesting a reduced repertoire of alternative routes for carbon metabolism (Bec et al. 2025). Under elevated CO_2_, *P. celeri* achieves doubling times of approximately two to three hours and areal productivities approaching 40 g m^−2^ day^−1^ in climate-simulated photobioreactors, while sustaining average productivities above 30 g m^−2^ day^−1^ over summer outdoor cultivation (Weissman et al. 2018; Krishnan et al. 2021). This high productivity, however, depends on supplemental CO_2_; under the cultivation conditions examined here, productivity declines by more than an order of magnitude when CO_2_-enriched air is replaced with ambient air. This pronounced dependence raises the possibility that the genome streamlining in *P. celeri* favors rapid growth under resource-rich conditions while limiting the metabolic flexibility needed to sustain growth under carbon limitation.

Rubisco catalyzes carbon fixation in the Calvin–Benson cycle, but its limited affinity for CO_2_ and competing oxygenase activity constrain photosynthesis when CO_2_ is scarce (Flamholz et al. 2019). Photosynthetic organisms have repeatedly evolved carbon-concentrating mechanisms (CCMs) that increase CO_2_ availability around Rubisco.

These include C_4_ photosynthesis, which arose independently in more than 60 plant lineages, and mechanisms based on carboxysomes in cyanobacteria and pyrenoids in eukaryotic algae (Sage et al. 2011; Hennacy and Jonikas 2020). However, an active CCM does not eliminate the need for photorespiration, as demonstrated in cyanobacteria and, more recently, during low-CO_2_ acclimation in *C. reinhardtii* (Eisenhut et al. 2008; Dao et al. 2025). Photorespiration detoxifies 2-phosphoglycolate by recycling its carbon into 3-phosphoglycerate for return to the Calvin–Benson cycle, at the cost of energy consumption and partial carbon loss as CO_2_ (Bauwe et al. 2010). In *P. celeri*, Ekness et al. (2026) found that productivity declined at higher pH and oxygen concentrations, although continued growth at high pH suggested a rudimentary CCM. These observations raise the question of whether its carbon-concentrating machinery operates efficiently enough to support rapid growth at low CO_2_, and how the metabolic demands of continued photorespiration contribute to the loss of productivity.

Elucidating complex responses such as acclimation to low CO_2_ in *P. celeri* requires a deeper understanding of how genotype gives rise to phenotype across multiple regulatory layers. Steichen et al. (2024) combined transcript profiling with metabolic flux analysis in *P. celeri* to identify a transcription factor related to the plant circadian regulators CCA1 and LHY that influences growth and carbon allocation under high light. A TG1 line expressing the TG2 version of this regulator showed a 15% increase in growth rate and a 25% increase in carbohydrate content. However, transcript abundance did not consistently track the corresponding metabolic fluxes. Integrating transcriptomics, proteomics, and metabolomics allows changes in gene expression to be compared with protein abundance and metabolite pools, although these measurements alone do not establish metabolic flux (Canelas et al. 2010).

Post-translational modifications provide an additional layer of regulation through which cells can alter the function of existing proteins. Phosphorylation, mediated by kinases and reversed by phosphatases, regulates enzyme activity, signal transduction, transcription, protein interactions, and subcellular localization. In *C. reinhardtii*, it controls the redistribution of light-harvesting complexes between photosystems, and specific phosphorylation events respond to CO_2_ limitation (Depège et al. 2003; Turkina et al. 2006). The functional consequences of most experimentally detected phosphosites remain unknown (Beltrao et al. 2012). Phosphosites are enriched in intrinsically disordered protein regions, whose relatively rapid sequence evolution contributes to the limited conservation of individual phosphorylation sites (Landry et al. 2009). Nevertheless, phosphosites with established functions tend to be more conserved than those without functional annotation, making evolutionary conservation a useful criterion for prioritizing candidate regulatory sites involved in acclimation to CO_2_ limitation (Landry et al. 2009).

Ubiquitination regulates protein fate and function through the covalent attachment of ubiquitin, typically to the ε-amino group of lysine residues. Proteins can carry individual ubiquitin molecules or chains whose linkage architecture influences their downstream effects. K48-linked chains are a major signal for proteasomal degradation, whereas monoubiquitination and K63-linked chains also regulate protein localization, DNA damage responses and signaling (Thrower et al. 2000; Hoege et al. 2002). Ubiquitin-binding receptors can also direct modified proteins to autophagic degradation (Pankiv et al. 2007). These functions make ubiquitination a potential contributor to CO_2_ acclimation through both selective protein turnover and regulation of retained proteins.

Lysine acetylation regulates chromatin organization and transcription, as well as the activity, interactions, and stability of non-histone proteins (Finkemeier et al. 2011). In *C. reinhardtii*, histone acetylation accompanies stress-responsive gene expression, while acetylation of carbon-metabolism proteins changes with light and acetate availability (Strenkert et al. 2011; Füßl et al. 2022).

CO_2_ can also regulate proteins directly through reversible carbamate formation on lysine ε-amino groups and protein N-termini. This reaction activates Rubisco through carbamylation of K201 (Lorimer 1981). However, carbamates readily dissociate during sample preparation, limiting their detection by conventional proteomics, whereas triethyloxonium-based trapping stabilizes native carbamates for mass spectrometry (Linthwaite et al. 2018). Cyanate reacts with lysine through its protonated form, isocyanic acid, to form the irreversible adduct homocitrulline (Wang et al. 2007).

LysCarComp-MS exploits competition between this reaction and reversible CO_2_ carbamylation to identify candidate CO_2_-binding sites (King et al. 2022). Cyanate can also arise from spontaneous decomposition of urea and carbamoyl phosphate, linking its production to nitrogen metabolism (Hagel et al. 1971; Guilloton and Karst 1987). In macrophages, integrated profiling identified shared targets of homocitrullination, acetylation, and phosphorylation in metabolic and ubiquitin-signaling pathways. Cyanate modification of diubiquitin also prevented its cleavage by the deubiquitinase OTULIN in vitro (You et al. 2023). The susceptibility of lysine ε-amino groups to ubiquitination, acetylation, reversible CO_2_ carbamylation, and irreversible homocitrullination makes these residues versatile sites for regulating enzyme activity and protein fate (Linthwaite et al. 2021; You et al. 2023). How these lysine modifications contribute to physiological responses to CO_2_ availability in photosynthetic eukaryotes remains poorly understood.

Paired transcriptome–proteome analyses have characterized algal nutrient responses (Schmollinger et al. 2014), and sequential enrichment has enabled joint profiling of the *Chlamydomonas* kinome and phosphoproteome (Werth et al. 2017), while combined ubiquitinome and proteome measurements have been applied to thermal adaptation in Phaeodactylum tricornutum (Li et al. 2026). Integrating these regulatory layers from the same biological samples under elevated and ambient CO_2_ allows us to develop a multiomics model of how different aspects of cell physiology contribute to the observed growth phenotype.

Here we integrate the transcriptome, proteome, phosphoproteome, ubiquitinome, acetylome, metabolome, and lipidome from the same biological replicates of *P. celeri* grown under elevated and ambient CO_2_ in climate-simulated photobioreactors. We complement these measurements with LysCarComp-MS profiling of candidate CO_2_-binding lysines and detection of homocitrulline in samples analyzed without cyanate treatment. Because CO_2_ supply also alters medium pH, the comparison captures the combined response to carbon limitation and alkalinization. Under ambient CO_2_, reduced investment in external nitrogen acquisition and cytosolic translation contrasts with preservation of plastid ribosomal proteins and increased abundance of carbon-fixation and photorespiratory enzymes. The switch between divergent Rubisco small-subunit isoforms points to holoenzyme remodeling as a potential mechanism for adjusting its catalytic properties to low CO_2_. These changes frequently oppose the corresponding transcript responses, placing post-transcriptional regulation at the center of the shift from biomass production toward cellular maintenance and carbon recovery.

Phosphorylation, ubiquitination, and acetylation provide additional routes through which metabolic state can influence protein function. We further propose that the similar chemistry of CO_2_ and cyanate enables them to compete directly for susceptible lysine ε-amino groups, linking metabolic activity to reversible carbamylation and persistent homocitrullination. Together, the phenotypic and multiomics data support an adaptation strategy in which *P. celeri* enters a quiescent state under low CO_2_, suspending growth while retaining the photosynthetic capacity needed to resume growth rapidly when CO_2_ is restored.

## Materials and methods Algal growth

### Strain and culture medium

*Picochlorum celeri* TG2 was isolated from Gulf of Mexico coastal water by high-light selection (Weissman et al. 2018; Becker et al. 2020). Cultures were grown in filter-sterilized Marine Dense medium (Weissman et al. 2018; Cano et al. 2024), a seawater medium of about 40 g L^−1^ Instant Ocean sea salt with urea (436 mg L^−1^) as the sole added nitrogen source, phosphate, an iron–EDTA chelate, trace metals and vitamins (Supplementary Methods 1).

Cell-free medium was sparged in the vessels under each gas for 24 h, the interval between daily dilutions, and its pH and total alkalinity were then measured (Ekness et al. 2026; Supplementary Methods 2). Medium sparged with air supplemented with 2.25% CO_2_ had a pH of 7.05 ± 0.10 and an alkalinity of 3.36 ± 0.14 meq L^−1^, whereas medium sparged with unsupplemented air had a pH of 8.23 ± 0.01 and an alkalinity of 2.19 ± 0.13 meq L^−1^ (mean ± SD of 12 titrations on four vessels), and a second medium batch measured months later agreed within 0.06 meq L^−1^ (Supplementary Data Set 1). A precipitate formed in the medium under air, but filtering it off before titration lowered the alkalinity by only 6%, which bounds the alkalinity held in the precipitate rather than in solution. With carbonate constants fitted to the dissolved-CO_2_ calibration of Ekness et al. (2026) for this medium (Supplementary Methods 2), the air-phase values correspond to about 450 ppm CO_2_ in the air stream, the level of ambient air, whereas the medium sparged with the nominal 2.25% stream held 250 to 280 µM dissolved CO_2_, equivalent to 1.4 to 1.5% CO_2_, which is the carbon status the cultures experienced within a dilution cycle. No external buffer was added, and dissolved inorganic carbon and pH changed together.

### Bioreactor set-up and operation

Cultures were grown in an automated ALGiSIM photobioreactor system (Cano et al. 2024), described in full by Karns (2024), in four square 500-mL Pyrex bottles (Corning, Corning, NY, USA) holding 400 mL of culture with a light path of about 8 cm (Weissman et al. 2018; Ekness et al. 2026), each illuminated on one face by cool-white LEDs, held at 33 °C by a thermoelectric module beneath it and mixed with a magnetic stir bar at about 125 rpm (Burch et al. 2025). The vessels shared a medium reservoir, deionized-water supply and gas stream (Figure 1A), culture volume was tracked by weight, and evaporative losses were replaced with deionized water every 2 h (Supplementary Methods 3).

**Figure 1.**
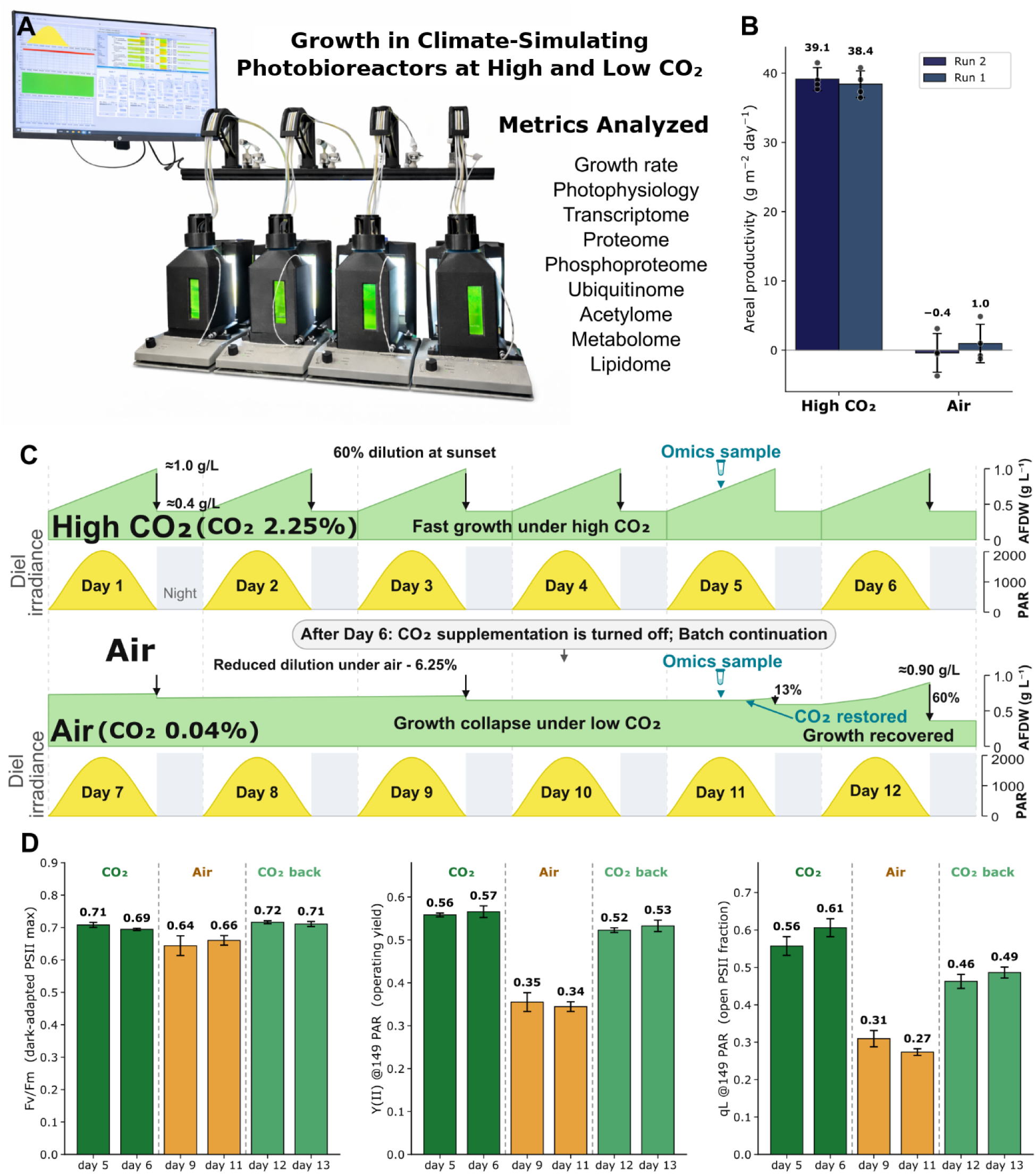
Growth arrest and recovery of *Picochlorum celeri* following removal and restoration of supplemental CO_2_ in climate-simulating photobioreactors. A) Climate-simulating photobioreactor system and the physiological and omics measurements collected during the experiment. Four parallel bioreactors were operated at 33 °C under a diel light regime reproducing a summer day in Mesa, Arizona (Supplementary Data Set 1). B) Areal biomass productivity under air supplemented with 2.25% CO_2_ (high CO_2_) and unsupplemented air in two independent runs. For each condition, productivity was averaged across the steady-state days within each bioreactor, and these per-bioreactor averages are shown as individual points. Bars show the mean ± SD across four bioreactors per run. High CO_2_ averages were calculated from four consecutive steady-state days under high CO_2_ in each run (Supplementary Data Set 1), whereas air averages used measurements from 2.5 d after CO_2_ removal onward. C) Schematic timeline showing biomass accumulation, diel irradiance, dilution, removal of supplemental CO_2_ after Day 6, the subsequent air phase, and restoration of CO_2_. Biomass for productivity measurements was sampled at sunset, whereas omics samples were collected at noon on Day 5 under high CO_2_ and on Day 11 in air. Dilution was reduced during the air phase as growth declined and restored after CO_2_ supplementation resumed. D) Photophysiology during the high CO_2_, air, and CO_2_-restored phases of run 2, shown as the dark-adapted maximum quantum yield of photosystem II (PSII), Fv/Fm, the operating quantum yield of PSII, Y(II), and the fraction of open PSII centers, qL. Y(II) and qL are shown at the 149 µmol photons m^−2^ s^−1^ step of the light curve. Bars show the daily mean ± SD of the four bioreactors sampled at noon.

The LED panels reproduced the photon-flux profile of a summer day in Mesa, Arizona (13 h light, 11 h dark, mid-day maximum 2,200 µmol photons m^−2^ s^−1^; Supplementary Data Set 1), calibrated in the culture bottle with a spherical quantum sensor (Heinz Walz GmbH; Supplementary Methods 3). Mass flow controllers (Aalborg Instruments and Controls, Orangeburg, NY, USA) blended CO_2_ into compressed house air to 2.25% (v/v), delivered at 400 mL min^−1^ per vessel, about one vessel volume per minute, through a sintered-glass sparger (Karns 2024; Burch et al. 2025; Supplementary Methods 3). CO_2_ limitation was imposed by closing the CO_2_ supply while maintaining air flow. At sunset, 60% of each culture was harvested and replaced with fresh medium.

Daily replacement was reduced to 10 to 15% during the air phase and increased to 60 to 65% during recovery (Supplementary Data Set 1).

Two wild-type runs of four vessels each were performed, with the days of the second run counted from June 21, 2026. In the first run, omics samples were collected from three vessels on Day 5 at high CO_2_ and on Day 11 in air, and biomass was sampled one, three and five days after CO_2_ withdrawal. In the second run, CO_2_ was withdrawn after Day 6 and restored on Day 11, with flow cytometry and fluorometry continued through Days 12 and 13.

Twelve additional samples came from four previously generated mutant lines of the alpha-class carbonic anhydrase genes PCCAA2 (CA3) and PCCAA1 (CAII) and the beta-class carbonic anhydrase genes PCCAB1 (CA4) and PCCAB2 (CA7), named as in Supplementary Data Set 2, made by Cas9 ribonucleoprotein editing as described (Burch et al. 2025; data not shown), and were included solely to estimate residual variance in the omics models. Each line was cultured in one vessel at 2.25% CO_2_ and sampled on three consecutive days.

### Sampling and ash-free dry weight

Biomass was sampled at the daily sunset harvest, and ash-free dry weight (AFDW) was measured in duplicate 5 to 20 mL aliquots on pre-ashed glass-fiber filters (TCLP, 0.7 µm, Pall, Port Washington, NY, USA). Filters were rinsed with 0.5 M ammonium formate, dried at 105 °C and weighed after cooling in a desiccator, then ashed at 550 °C and weighed again (Weissman et al. 2018; Ekness et al. 2026). Areal productivity was the volumetric productivity multiplied by 67 L m^−2^, the 0.40 L culture volume divided by the 0.006 m^2^ illuminated face of the bottle (Weissman et al. 2018).

Omics samples were collected at simulated solar noon as 200 mL of culture per vessel, split into four 50-mL conical tubes, which were centrifuged for 5 min at 3,000 × g and 4 °C, resuspended in 20 mL of deionized water each, centrifuged again and drained. The pellets were frozen in liquid nitrogen, sent on dry ice to the Environmental Molecular Sciences Laboratory (EMSL, Pacific Northwest National Laboratory, Richland, WA, USA; project 61507) and lyophilized. In the second run, all four vessels were sampled at simulated solar noon for flow cytometry and fluorometry.

### Phenotypic analysis Photophysiology

Chlorophyll fluorescence was recorded with a Dual-PAM-100 (Heinz Walz GmbH, Effeltrich, Germany). Samples taken at simulated solar noon were dark-adapted for 20 min without far-red pre-illumination, and Fo and Fm were determined with a saturating pulse before light curves consisting of 50-s actinic steps from 6 to 3,242 µmol photons m^−2^ s^−1^, which describe induction rather than steady-state responses. Y(II) and qL were evaluated at the 149 µmol photons m^−2^ s^−1^ step. Fv/Fm describes maximum PSII photochemical yield, Y(II) operating yield, and qL the fraction of open PSII centers under the lake model (Kramer et al. 2004; Baker 2008).

Daily means over the four vessels were compared between phases by Welch’s t-test, Days 5 and 6 under CO_2_ against Days 9 and 11 in air, and Days 12 and 13 after CO_2_ restoration are reported as recovery without a test. P700 measurements were excluded because the dark-adapted maximum could not be determined reliably.

### Flow cytometry and cell size

Cells were diluted 1,000-fold in deionized water and analyzed immediately on an Attune NxT flow cytometer (Thermo Fisher Scientific, Waltham, MA, USA) with a 488 nm laser, drawing 200 µL at 200 µL min^−1^ so that the event rate stayed near 1,000 events s^−1^, which keeps counting efficiency consistent between samples, recording forward scatter (FSC, 360 V), side scatter (SSC, 300 V) and chlorophyll autofluorescence (BL3, 695/40 nm, 340 V) with an FSC-H acquisition threshold of 23,000 and an area scaling factor of 1.04. Files were read with FlowKit 1.3.2, cells were events with FSC-A above 20,000 and SSC-A above 2,000 and below saturation, the pulse-shape gate retained events with FSC-A − 1.0259 × FSC-H ≤ 22,204, and per-sample chlorophyll autofluorescence was the median BL3-A of events retained by the pulse-shape gate, with low-chlorophyll cells defined as events retained by the pulse-shape gate with BL3-A below 1,000 (Supplementary Methods 4).

Cell diameter was obtained from the median FSC-A of events retained by the pulse-shape gate, using a Mie-theory forward-scatter model calibrated against 2, 4 and 6 µm polystyrene beads at a fixed cell refractive index of 1.41 (Supplementary Methods 4). Daily values are the mean and standard deviation across the retained vessels (Supplementary Data Set 1 and Supplementary Figure 1).

### Protein content

Total protein per cell was measured by amino acid analysis at the Molecular Structure Facility, University of California, Davis (Hitachi LA8080 analyzer with post-column ninhydrin detection; Hitachi High-Tech, Tokyo, Japan). Aliquots of 2 × 10^8^ cells, counted at the bench with the cytometer’s own software, from each of the four vessels of the second run, taken on Day 6 under CO_2_ and Day 11 in air, were hydrolyzed in 6 N HCl with 1% phenol at 110 °C for 24 h with norleucine as internal standard (Supplementary Methods 5). Protein per cell was the summed residue mass of the quantified amino acids, uncorrected for hydrolysis losses of tryptophan, serine, threonine and cysteine, and was expressed per gram of AFDW with the same-day AFDW per cell of the same vessel. Conditions were compared by Welch’s t-test on the four vessels.

### Transcriptomics

#### RNA extraction, library preparation and sequencing

Total RNA was extracted from the pellets of the three wild-type vessels under each condition and the 12 additional samples with the Quick-RNA Miniprep kit (Zymo Research; R1055), and integrity was assessed on a TapeStation (Agilent Technologies, Santa Clara, CA, USA) with an RNA integrity number above 7 required. Strand-specific poly(A) libraries (Illumina Stranded mRNA Prep with 10-bp unique dual indexes; Illumina, San Diego, CA, USA) were sequenced by SeqCenter (Pittsburgh, PA, USA) on an Illumina NovaSeq X Plus as 150-bp paired-end reads to 30 million pairs per library.

Organellar genes were excluded from the transcriptome tests because the libraries were poly(A)-selected.

### Read processing and differential expression

Reads were trimmed with fastp 1.0.1 (adapter detection, overlap-based correction, sliding-window quality trimming, minimum length 36 nt), which retained 99.2% of read pairs (Supplementary Methods 12). They were aligned with STAR 2.7.11b in two-pass mode to the diploid *P. celeri* assembly (15 chromosome pairs; Becker et al. 2020) with the gene models of this study (13,746 nuclear genes plus the chloroplast and mitochondrial genomes; Supplementary Methods 12), retaining reads from both haplotypes, and strandedness and transcript integrity were confirmed with RSeQC

5.0.3. Fragments were counted per transcript with featureCounts 2.1.1, reverse-stranded and with multi-mapping fragments counted once, and allele-level counts were summed into gene-level counts through the 6,351 allelic pairs defined by nucmer alignment of the two haplotypes (Supplementary Methods 12). Genes were tested with DESeq2 1.42.0 (Love et al. 2014) in the six-group model with the contrast wild type in air against wild type at high CO_2_, apeglm shrinkage of the fold change and Benjamini– Hochberg (BH) adjustment (see Statistical analysis). Of 7,352 tested genes, those with adjusted p below 0.05 and an absolute shrunken log_2_ fold change of at least 1 were counted as differentially expressed.

### Curated pathways and enrichment

Because KEGG pathway assignments are sparse for a non-model alga, genes were grouped into 44 pathway sets defined for this study (Supplementary Data Set 3), and set-level changes were tested with fry (limma; Wu et al. 2010; Ritchie et al. 2015) and camera on log_2_ counts per million with the same design and contrast. KEGG pathway and Gene Ontology enrichment, with human-disease and non-plant terms removed, was run as over-representation analysis with clusterProfiler 4.10.0 against the 7,352 tested genes and as gene-set enrichment with fgsea 1.28.0 on genes ranked by the DESeq2 Wald statistic, with BH adjustment within each test and collection (Supplementary Methods 12; Supplementary Data Set 3).

### Proteomics

### Extraction, digestion and TMT labeling

From each lyophilized pellet, 10 mg underwent MPLEx extraction at EMSL (Nakayasu et al. 2016), whose aqueous and organic layers were retained for metabolomics and lipidomics, with three process blanks. The remaining material (about 50 mg) was lysed in SDS with phosphatase, protease, deacetylase and deubiquitinase inhibitors, and about 2 mg of protein per sample was reduced, alkylated, digested with trypsin and Lys-C on S-Trap columns and desalted (Supplementary Methods 6). Diglycine-remnant (K-ε-GG) peptides were enriched from 430 µg of each digest, and 200 µg of the depleted peptides was labeled with TMTpro 18-plex reagent. The plex held the six wild-type samples and the 12 additional samples with no pooled reference channel (Supplementary Data Set 2), and the pooled peptides were fractionated by high-pH reversed-phase chromatography into 12 concatenated fractions, of which 5% went to global proteomics and 95% to Fe(III)-NTA phosphopeptide enrichment (Supplementary Methods 6).

### Liquid chromatography and mass spectrometry

Global, K-ε-GG and acetyl-lysine samples were separated on a nanoAcquity UPLC (Waters, Milford, MA, USA) and analyzed on an Orbitrap Eclipse Tribrid (Thermo Fisher Scientific) with a FAIMS Pro interface, and phosphopeptides were separated on a Vanquish Neo and analyzed on an Orbitrap Exploris 480 without FAIMS, both by data-dependent acquisition with higher-energy collisional dissociation (HCD; Supplementary Methods 7).

### Peptide identification and protein quantification

Spectra were searched with FragPipe 24.0 (MSFragger 4.4; Kong et al. 2017) against the non-redundant *P. celeri* proteome (7,575 sequences, one allele per allelic pair, the unpaired alleles of both haplotypes and the 87 organellar proteins) with common contaminants except human ubiquitin and NEDD8 (Supplementary Methods 7) and reversed decoys, with carbamidomethyl cysteine and TMTpro on lysine fixed, and

TMTpro on peptide N-termini, methionine oxidation and protein N-terminal acetylation variable (Supplementary Methods 7). Peptide-spectrum matches (PSMs) were rescored with Percolator and filtered with Philosopher to 1% false discovery rate (FDR) at the PSM and protein levels, and reporter-ion intensities were extracted with TMT-Integrator (PSM probability ≥ 0.9, precursor purity ≥ 0.5, virtual reference channel). Proteins sharing all their peptides were merged into groups and quantified with MSstatsTMT 2.10.0 (Huang et al. 2020) from group-unique peptides with per-channel median normalization and no reference-channel normalization, in the six-group model with the contrast wild type in air minus wild type at high CO_2_ and BH adjustment. Transcript– protein fold-change correlations used finite estimates for nuclear genes with a unique transcript and a single unambiguously mapped protein group. Of the 6,259 quantified protein groups, those with adjusted p below 0.05 and an absolute log_2_ fold change of at least 0.5 were counted as differentially abundant. Set tests were run as for the transcriptome on the 6,199 protein groups summarized in all 18 channels, a universe that includes the chloroplast-encoded proteins, with the KEGG and GO gene-set enrichment ranked on the moderated t statistic of a limma fit to the same summarized abundances.

### Absolute protein abundance and proteome allocation

Relative protein abundance was computed by the total protein approach (Wiśniewski and Rakus 2014) as the integrated MS1 precursor area of each protein group, apportioned to channels by the TMT reporter fractions and summed, expressed as a percentage of the summed area of all quantified groups in the same channel. Proteome allocation is the share of total protein mass in each curated category, drawn as a squarified treemap. Amounts per cell and per gram of biomass (Supplementary Data Set 2) were obtained by scaling these mass fractions to the total protein per cell measured by amino acid analysis, 1.76 and 1.22 pg for CO_2_ and air. Measured AFDW per cell (4.76 and 2.49 pg for CO_2_ and air) was used for biomass normalization, with ash assumed at 15% of dry weight (Supplementary Methods 5). Measured cell diameters (2.52 and 2.30 µm for CO_2_ and air) were used to derive cell volumes for molar concentrations. All measured physiological constants were obtained in the second run and applied to the proteomics samples of the first run. Dividing the same precursor areas by the count of distinct theoretical fully tryptic peptides of 7 to 50 residues, which is intensity-based absolute quantification (Schwanhäusser et al. 2011), changes the median group by 0.5% and moves 87 of 6,192 groups by more than twofold, and those columns are retained in Supplementary Data Set 2.

### Phosphoproteomics

Phosphopeptide spectra were searched with the same FragPipe workflow as the global proteome, except that TMTpro was fixed on peptide N-termini as well as on lysine, phosphorylation of serine, threonine and tyrosine was allowed as a variable modification (up to three per peptide, four variable modifications in total) and the precursor tolerance was 10 ppm (Supplementary Methods 7). Phosphosites were localized with PTMProphet, and reporter-ion intensities were extracted with TMT-Integrator for sites with a localization probability of at least 0.75, a PSM probability of at least 0.5 and a precursor purity of at least 0.5. MSstatsPTM 2.4.1 (Kohler et al. 2023) then summarized the phosphosite and global-proteome runs separately from PSMs with a localization probability of at least 0.75, a precursor purity of at least 0.6 and a PSM probability of at least 0.7, using unique peptides, median normalization within each run and model-based imputation, and tested the wild-type contrast in the six-group model, which left 29,076 sites after removal of 13 mapping to common laboratory contaminants (porcine trypsin and human keratins). The protein-adjusted change of each site was its log_2_ fold change minus κ times the protein log_2_ fold change, with κ = 0.46 for phosphorylation from passenger-peptide calibration, the variance propagated from both standard errors, a t distribution on Welch–Satterthwaite degrees of freedom and BH adjustment at 0.05 within the reportable sites (Supplementary Methods 8). Set tests used fry on per-channel protein-adjusted site abundances, with sites rather than genes as set members and κ = 0.469 from the passenger-peptide regression (Supplementary Methods 8). Kinase-recognition contexts from ±7-residue windows were tested for directional enrichment by Fisher’s exact test, and phosphosite annotations from EPSD, PhosPhAt and UniProtKB/Swiss-Prot were transferred by ortholog alignment (Supplementary Data Set 4).

### Ubiquitinomics

K-ε-GG peptides were enriched from 430 µg of each digest with anti-K-ε-GG antibody beads (PTMScan HS Ubiquitin/SUMO Remnant Motif kit, Cell Signaling Technology, Danvers, MA, USA), labeled on the beads with TMTpro, pooled and analyzed unfractionated in two acquisitions (Supplementary Methods 9). Spectra were searched as for the global proteome but with a 10-ppm precursor tolerance, up to three missed cleavages, TMTpro fixed on peptide N-termini only, and diglycine on lysine, TMTpro on lysine, methionine oxidation and protein N-terminal acetylation as variable modifications (at most four per peptide), giving 14,024 PSMs on 2,340 proteins at 1% PSM and protein FDR (Supplementary Methods 7). Sites were localized as for the phosphosites and summarized and modeled with MSstatsPTM, with model-based imputation enabled for both the PTM and global-protein data. This yielded 4,772 sites, of which 3,498 had finite fold-change estimates for the quantified arm. Because quantified-arm fold changes are relative to the channel median, the global loss of sites in air was read from raw reporter signal and detection rates. Each site was corrected for its protein with κ = 0.736 and called changed when it passed BH at 0.05 (Supplementary Methods 8). Sites detected in at least two of three replicates of one condition and in none of the other formed the binary arm. Detection calls used only observed, non-censored features, excluding imputed intensities. The binary censoring test likewise used observed feature intensities and protein-change estimates from non-imputed data. Because a site also disappears when its protein falls below the detection floor, each such site was tested on the raw reporter scale for whether the intensity expected from the protein change would still have been detectable, and was called lost or gained when it would, explained by the protein change when it would not, and indeterminate when the expected intensity lay within the uncertainty of the floor (Supplementary Methods 9; Supplementary Data Set 4). Chain-linkage abundance was read from K-ε-GG PSMs on the seven chain-forming lysines of ubiquitin, summed per linkage and channel and compared between conditions and against the pool of all other sites (Supplementary Methods 9; Supplementary Data Set 4). The quantified arm was tested against the curated sets with fry and the binary arm with a one-sided Fisher’s exact test against the proteins carrying a measured site, with BH within each column. Site conservation was assessed against the *Arabidopsis thaliana* ubiquitinome of Song et al. (2024) through reciprocal-best-hit orthologs with the Mantel–Haenszel test conditional on protein, and the ubiquitin–proteasome system was censused from the InterPro and Pfam domains of the annotation (Supplementary Data Set 4).

### Lysine homocitrulline and acetyl-lysine sites Cyanate competition and acetyl-lysine enrichment

Cell pellets from the same 18 samples were lysed in 50 mM HEPES pH 7.2 with protease and deacetylase inhibitors and quantified by BCA (Supplementary Methods 10). Each lysate was split into two 50 µL reactions in degassed 200 mM potassium phosphate pH 7.2 with 50 mM potassium cyanate, one receiving 50 mM NaHCO_3_ and the other 50 mM NaCl, and sealed against gas exchange for 1 h at about 25 °C. In this LysCarComp-MS design (King et al. 2022), changes in cyanate-dependent lysine labeling as homocitrulline were measured in the presence versus absence of added bicarbonate. Proteins were precipitated, digested with trypsin and labeled with TMTpro 18-plex in two plexes, each holding both halves of nine lysates (Supplementary Data Set 2), which were fractionated by high-pH reversed-phase chromatography into 4 pooled fractions per plex, enriched for acetyl-lysine peptides with antibody beads that also bind homocitrulline (PTMScan HS Acetyl-Lysine Motif kit, Cell Signaling Technology; Martinez-Val et al. 2017), and analyzed on the Orbitrap Eclipse as above (Supplementary Methods 10).

### Database searches and site quantification

Spectra were searched as for the ubiquitinome except that diglycine was replaced by acetyl-lysine (+42.0106 Da) and carbamyl-lysine (+43.0058 Da, homocitrulline) as variable modifications (at most five per peptide), and TMT-Integrator produced separate site tables for the two tags (Supplementary Methods 7 and 11). Acetyl-lysine sites were quantified with MSstatsPTM as above from the 18 control-half channels (cyanate with NaCl), with the protein term from the unenriched global proteome of the same cultures, κ = 0.727 (Supplementary Methods 8), BH adjustment over the 3,804 sites with a finite estimate and standard error (of 4,654) and fry set tests on per-channel adjusted site abundances averaged per gene. For the bicarbonate competition, the ratio of each homocitrulline site was the log_2_ of its reporter intensity in the NaCl half over the NaHCO_3_ half of the same lysate, centered on the median of all homocitrulline sites of that lysate because homocitrulline and acetyl-lysine peptides compete for the same antibody (Supplementary Methods 10). Sites quantified in at least 12 of the 18 lysates (1,361 of 2,777) were tested for a nonzero grand-mean ratio with limma, with plex as a covariate, empirical-Bayes moderation and BH at 0.05, positive deviations being called protected and negative ones exposed, and candidates in the sense of King et al. (2022) additionally exceeded the population mean by two standard deviations on an internal lysine. Sequence context, protein abundance and set membership were compared between the two classes (Supplementary Methods 10; Supplementary Data Set 4).

### Homocitrulline in the unenriched proteome

The global, phosphopeptide-enriched and K-ε-GG-enriched data were re-searched with TMTpro on lysine, carbamyl-lysine and acetyl-lysine as variable modifications, carbamylation of protein N-termini being allowed in the global search, with PSMs validated by Percolator 3.7.1 and filtered to 1% picked protein FDR (Supplementary Methods 10). Homocitrulline matches were kept when the observed mass lay within 0.02 Da of the assigned one and the modified lysine was internal to the peptide, because trypsin does not cleave after homocitrulline and a C-terminal carbamyl-lysine must have formed after digestion. Within each channel, homocitrulline reporter intensities were divided by those of unmodified matches from the same proteins, and the three vessels per condition were compared by Welch’s t-test. For the 35 sites on 25 proteins quantified in all six wild-type channels, the site log_2_ ratio was regressed on the MSstatsTMT protein log_2_ fold change with a case bootstrap over proteins and leave-one-protein-out sensitivity analyses (Supplementary Methods 10; Supplementary Data Set 4; Supplementary Figure 2). Acetylation-site conservation against *Chlamydomonas reinhardtii* (Füßl et al. 2022) was tested through reciprocal-best-hit orthologs with the Cochran–Mantel–Haenszel test stratified by protein pair (Supplementary Methods 10).

### Metabolomics

The aqueous MPLEx extracts were analyzed by GC-MS after methoximation and trimethylsilylation on an Agilent 7890A with a 5975C detector, with spectra deconvoluted in Metabolite Detector and matched against the PNNL-augmented Agilent library, NIST20 and Wiley11, and by HILIC LC-MS/MS on an Orbitrap Eclipse Tribrid in negative mode, with features aligned in Compound Discoverer 3.3 (0.5-min retention-time shift, 5-ppm mass tolerance, minimum intensity 2.5 × 10^5^; Supplementary Methods 11). Identifications were manually validated at the facility’s four levels, level 1 (MS1, MS2 and retention-time match), level 2 (MS2 match), level 3 (MS1 and retention-time match) and level 4 (MS1 with a partial MS2 match), and the retained set held 83 level-1, 14 level-2 and two level-3 metabolites. HILIC features below three times the process blank were excluded, metabolites detected on both platforms were taken from the platform with the higher within-platform intensity percentile, and intensities underwent probabilistic quotient normalization (PQN; Dieterle et al. 2006) within each platform across the 18 samples. The 99 metabolites were tested with limma in the six-group model with empirical-Bayes moderation (trend and robust options) and BH adjustment. Because PQN set most of the significance calls, sensitivity analyses with unnormalized intensities and wild-type samples alone are reported in Supplementary Methods 11 and Supplementary Data Set 2. Category tests used fry on 12 enzyme-derived metabolite sets, excluding four metabolites with opposite directions across platforms, and ATP, GTP, NADH, NADPH and acetyl-CoA were not recovered.

### Lipidomics

The organic MPLEx extracts were dried, resuspended in 200 µL of 10% chloroform in methanol and separated by reversed-phase chromatography on a CSH C18 column over a 21-min gradient, with detection on an Orbitrap Fusion Lumos in separate positive- and negative-mode runs (Supplementary Methods 11). Lipids were identified with LIQUID from the diagnostic head-group and acyl-chain fragments in the tandem spectra, with the precursor mass error, isotopic profile and extracted-ion chromatogram examined for every identification. The identified lipids, with their retention times and observed m/z, formed a target list against which MZmine 2 aligned and gap-filled the features of every run, and all aligned features were manually verified, giving 307 lipid names across 16 subclasses. Fourteen positive-mode pairs of positional isomers shared one chromatographic peak, with identical intensities in every wild-type sample, QC pool and blank, and each pair was counted once under a joined name, giving 271 measured features. Species with pooled-QC relative standard deviation above 30% (32 of 271) or fewer than two finite wild-type values per condition were excluded, leaving 237 species. No internal standards were included, and all lipid values are therefore relative peak intensities rather than absolute amounts. Because extracts represented equal biomass, log_2_ peak intensities were tested without sample normalization in the metabolomics limma model, with PQN as a sensitivity analysis (Supplementary Methods 11). The double-bond index was the signal-weighted number of double bonds per acyl chain, class-level shifts were tested by Wilcoxon signed-rank test across species within each class, and species of the four thylakoid classes were compared with all other species by Mann–Whitney test on log_2_ fold changes. Because ionization efficiency differs between lipid classes, only fold changes were compared and no share of total signal was computed. Lipid-metabolic enzymes were grouped by EC-defined pathway step, and desaturation was compared with the other steps by Mann–Whitney test.

### Statistical analysis

All omics layers were tested in one design in which each of the 18 samples belongs to one of six groups, wild type at high CO_2_, wild type in air and the four carbonic anhydrase lines at high CO_2_, and every layer was fitted with a model carrying one coefficient per group and the model appropriate to its data type (a negative-binomial generalized linear model in DESeq2 for transcript counts, linear models on log_2_ intensities in MSstatsTMT for protein groups, MSstatsPTM for modification sites and limma for metabolites and lipids), with the single contrast wild type in air minus wild type at high CO_2_. The four mutant groups enter no contrast and only supply residual variance, which raises the residual degrees of freedom of the contrast from 4 to 12 without changing its estimate.

Residuals were treated as independent without a term for the vessel, and the wild-type comparison therefore rests on three vessels per condition. Fold changes are air relative to high CO_2_, and p-values were adjusted by the BH method within each layer and, for set tests, within each collection. The bicarbonate competition compared the two halves of each lysate in a grand-mean model over the 18 lysates with plex as a covariate, whereas the measurements of the second bioreactor run compare phases within the same four vessels by the test stated in each legend, a phase effect describing the combined change in gas supply, pH and dilution rate. Principal component analysis (PCA), proteome allocation and genome-position plots used the six wild-type samples only, and all analyses were run in R 4.3.3 and Python 3.10 to 3.12 with the packages named above. Public archiving of the analysis scripts, which call the packages named above, is pending.

### Accession numbers and data availability

Processed results are provided in Supplementary Data Sets 1 to 4, with sample and channel assignments, acquisition filenames and reference-file checksums in Supplementary Data Set 2. Complete sample-level model inputs, including all 18 samples or paired lysates, exact reference sequences, annotation files and the original PAM export are supplied in Supplementary File 1. Released mass spectrometry data are available through EMSL Science Central under project 61507 (https://sc-data.emsl.pnnl.gov/?projectId=61507). The file-level map in Supplementary Data Set 2 distinguishes acquisitions with established EMSL dataset identifiers from entries without them. Deposition of raw sequencing reads in the NCBI Sequence Read Archive, proteomics data in PRIDE, the remaining physiological source data and selected analysis scripts remains pending, with Zenodo planned for the code. Repository accessions and a Zenodo DOI will be added when available.

## Supplementary data

Supplementary Data Set 1. Cultivation and physiology.

Supplementary Data Set 2. Omics measurements and differential analyses.

Supplementary Data Set 3. Gene sets and enrichment.

Supplementary Data Set 4. PTM site analyses.

Supplementary File 1. Complete model inputs, reference sequences, annotations and original PAM measurements.

Supplementary Information. Supplementary Methods 1 to 12, references and Supplementary Figures 1 and 2.

## Results

### CO_2_ limitation arrests growth reversibly while photosynthetic capacity is retained

We grew wild-type *P. celeri* in climate-simulating photobioreactors (Figure 1A) that reproduce the light of a summer day (June 30^th^) in Mesa, Arizona, at 33 °C, in four identical vessels that receive the same medium and the same gas. The vessels were sparged either with air supplemented to 2.25% CO_2_, the condition under which this alga reaches its published growth rates, or with unsupplemented air. In this medium CO_2_ sets the pH, and the air condition was therefore also the more alkaline one, near pH 8.2 against pH 7 under supplemented air (Ekness et al. 2026). We deliberately left the medium unbuffered because an open pond experiences the same coupling, and the air condition throughout this work therefore combines low CO_2_ with high pH.

Under high CO_2_, cultures achieved areal productivities of approximately 39 g m^−2^ d^−1^ in both experimental runs. Following transfer to air, growth persisted for the first 24 h at roughly two thirds of this rate before declining to near zero, where it remained for the duration of the low-CO_2_ treatment (Figure 1B, Supplementary Data Set 1). To determine whether the cultures remained viable after this prolonged period without growth, we repeated the experiment and restored CO_2_ after five days in air (Figure 1C). Biomass accumulation was measured within three hours of CO_2_ restoration, and over the following day the cultures reached a productivity of 43 g m^−2^ d^−1^, comparable to that observed before CO_2_ withdrawal.

The rapid recovery of growth indicated that the cells had not been irreversibly damaged but instead remained in a quiescent, growth-ready state. To determine whether the photosynthetic capacity was similarly preserved during carbon limitation, we monitored the cultures using pulse-amplitude-modulated fluorometry (Figure 1D). The maximum quantum yield of photosystem II photochemistry in dark-adapted cells, Fv/Fm, is widely used as an indicator of photosystem health (Baker 2008). Fv/Fm was only 7% lower in air than under high CO_2_, a reduction within the diel variation observed under high CO_2_, and recovered fully within one day of CO_2_ restoration. Light-response curves collected after each dark-adapted measurement were used to determine Y(II), the operating quantum yield of photosystem II, and qL, the fraction of photosystem II reaction centers remaining open, at the 149 µmol photons m^−2^ s^−1^ actinic step (Kramer et al. 2004). Both parameters were substantially more affected than Fv/Fm; Y(II) declined by approximately one third in air, while qL declined by half. Following CO_2_ restoration, Y(II) recovered to approximately 95% of its high-CO_2_ value within one day, whereas qL reached only approximately 80% after two days.

Flow cytometry showed that cells became smaller under air, with diameter decreasing from 2.5 to 2.3 µm over the first three days, corresponding to a 25% reduction in cell volume (Supplementary Data Set 1). Protein per cell decreased by 31%, roughly matching the reduction in cell volume, so protein concentration per unit cell volume remained essentially unchanged. Chlorophyll autofluorescence per cell similarly declined by 24%, indicating that chlorophyll autofluorescence was largely maintained relative to cell volume during carbon limitation. In contrast, ash-free dry mass per cell decreased by 48%, indicating a disproportionate loss of non-protein biomass as growth ceased. Following CO_2_ restoration, cell size and chlorophyll autofluorescence recovered rapidly.

Taken together, these observations indicate that the air treatment induces a reversible growth arrest without a corresponding loss of photosynthetic capacity. The near-zero productivity and disproportionate depletion of non-protein biomass are consistent with the possibility that internal carbon reserves are consumed to support maintenance metabolism once growth ceases. At the same time, the fluorescence response is more consistent with restricted electron use downstream of PSII than with substantial photodamage (Baker 2008). Thus, although the air treatment combines low CO_2_ with increased pH, the growth arrest appears to reflect insufficient carbon availability to sustain carbon fixation and associated electron consumption rather than deterioration of the photosynthetic apparatus.

### Dissection of the transcriptome and proteome of *P. celeri* under high and low CO_2_ conditions

CO_2_ limitation was accompanied by extensive changes in both the transcriptome and proteome of *P. celeri*. Air-grown and high-CO_2_ samples separated clearly along PC1, which explained 95.2% of transcriptome variance and 89.8% of proteome variance (Figures 2A and 2C). In air, 1,358 transcripts increased and 1,041 decreased among the 7,352 measured (adjusted *P* < 0.05; |log_2_FC| ≥ 1), while 1,148 proteins became more abundant and 905 less abundant among the 6,259 quantified protein groups (adjusted *P* < 0.05; |log_2_FC| ≥ 0.5; Figures 2B and 2D). Considering statistical significance alone, 62.8% of transcripts and 70.8% of proteins differed between conditions, revealing extensive remodeling of both the transcriptome and proteome.

**Figure 2.**
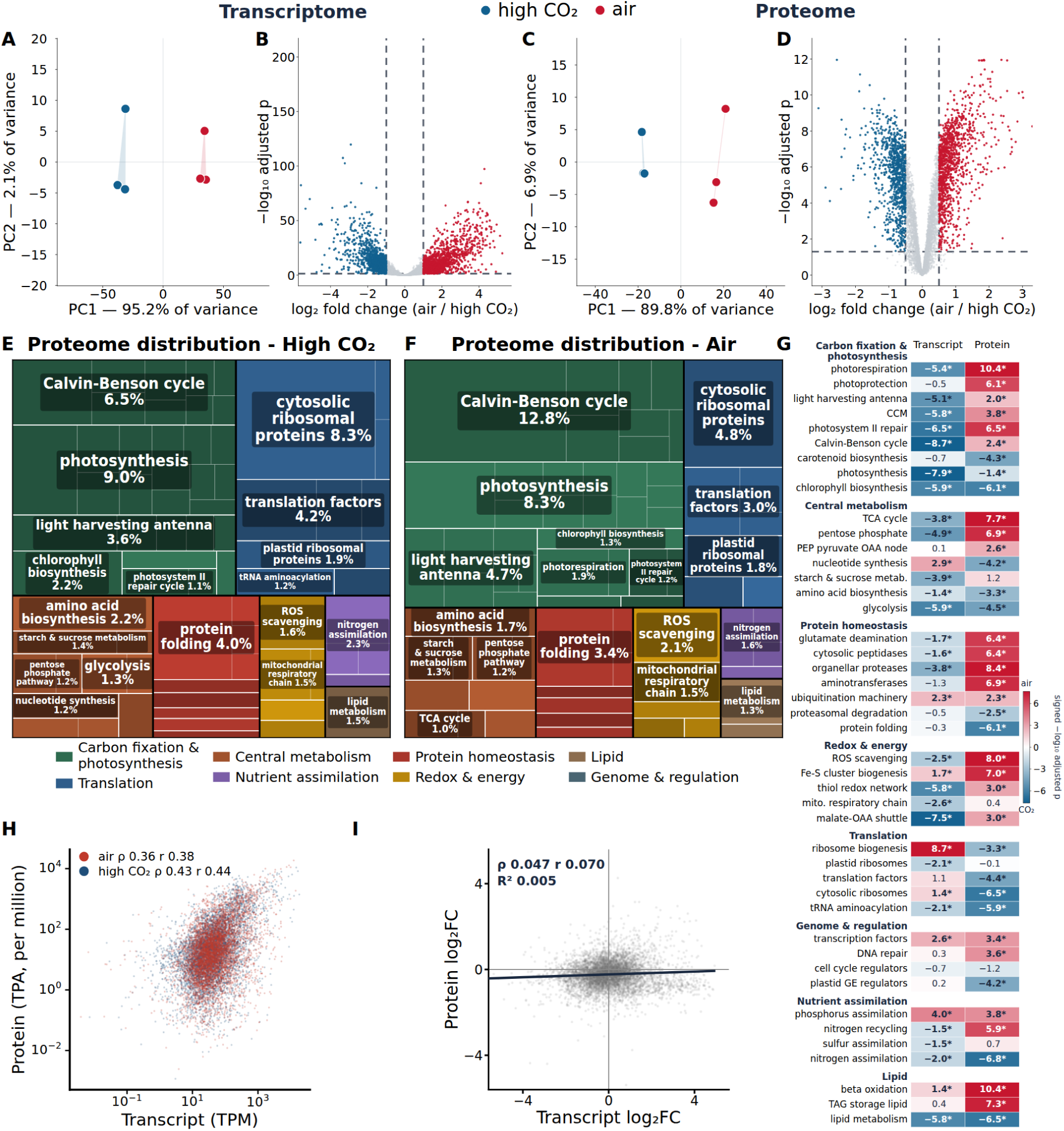
Transcriptome and proteome remodeling under CO_2_ limitation. Blue denotes high CO_2_ and red denotes air throughout. A, C) Principal-component analysis of transcript (A) and protein (C) abundance under high CO_2_ and air, with each point one bioreactor (n = 3 per condition). Axes indicate the proportion of total variance explained. B, D) Transcript (B) and protein (D) responses to air relative to high CO_2_, with positive values indicating higher abundance in air. Features meeting adjusted p < 0.05 and |log_2_ fold change| ≥ 1 for transcripts or ≥ 0.5 for proteins are colored red when higher in air and blue when higher under high CO_2_, and the remaining features are gray. E, F) Proteome investment under high CO_2_ (E) and air (F). Tile area is the protein-mass fraction from the total protein approach assigned to each curated cellular process within the displayed annotated subset, which excludes the 34% of quantified protein mass not assigned to a curated process, with the same category colors in both conditions. Labels show percentages of the total quantified proteome, averaged across the three wild-type (WT) channels per condition, and are omitted from tiles too small to carry them, including the smallest category (genome and regulation, 0.5% of the proteome). G) Transcript- and protein-level responses of the 44 curated pathway sets (fry test). Cell color shows the signed −log_10_ adjusted p, red for sets with higher abundance in air and blue for sets with higher abundance under high CO_2_, saturating at ±6. Printed values are the signed −log_10_ adjusted p, and asterisks mark adjusted p < 0.05. H) Relationship between condition-mean transcript and protein abundance under air and high CO_2_ across 5,911 unambiguously paired nuclear genes. Transcript abundance is mean transcripts per million (TPM), and protein abundance is the total-protein-approach mass fraction divided by molecular weight, normalized within each channel to one million and averaged across three WT replicates per condition. Spearman (ρ) and Pearson (r, on log_10_ values) correlations are calculated from the condition means. I) Relationship between transcript and protein log_2_ fold changes for air relative to high CO_2_ across 5,982 unambiguously paired nuclear genes. ρ is Spearman’s and r Pearson’s correlation, and the line is the ordinary least-squares fit of protein on transcript log_2_ fold change with its R^2^.

To identify the processes most affected by this remodeling, we grouped genes into 44 functional categories for proteome allocation and gene-set enrichment analyses (Supporting Information). The photosynthetic core, comprising the photosystem cores, oxygen-evolving complex, electron-transfer proteins, and chloroplast ATP synthase, accounted for 9.0% of total protein mass under high CO_2_ and declined to 8.3% in air (Figures 2E and 2F). This modest decline concealed distinct responses within the photosynthetic machinery. Allocation to the four major PSII core proteins, D1, D2, CP43 and CP47, remained nearly constant, at 1.71% under high CO_2_ and 1.76% in air, whereas the proteome share allocated to chloroplast ATP synthase declined by approximately 33%, from 2.68% to 1.80%, with the α and β subunits contributing to this reduction. By contrast, the separately classified light-harvesting antenna expanded from 3.6% to 4.7%, with increases distributed across the chlorophyll a/b-binding protein family. Allocation to photoprotection also increased, rising approximately twofold from 0.38% to 0.71% of the proteome, driven largely by PsbS.

Allocation to the broader carbon fixation and photosynthesis group increased from 24.0% of total protein mass under high CO_2_ to 31.3% in air, largely through expansion of the Calvin–Benson cycle, whose proteome share doubled from 6.5% to 12.8%. The Rubisco large subunit accounted for most of this increase, rising from 2.33% to 7.39% of the proteome, while the major Rubisco activase increased from 0.35% to 1.31%. This greater allocation to Rubisco was accompanied by a switch in small-subunit composition. *P. celeri* carries three *RbcS* genes that encode two divergent mature isoforms sharing 69.9% amino acid identity: RbcS1/2, encoded by two genes, and RbcS5, encoded by one. The proteome share of RbcS1/2 decreased from 0.46% to 0.14%, whereas RbcS5 increased from 0.08% to 1.10%, reversing their relative contributions while increasing their combined proteome share approximately 2.3-fold, from 0.54% to 1.24% (Supplementary Data Set 2). Alongside these changes in carbon fixation, allocation to photorespiration increased from 0.74% to 1.94%, while allocation to the carbon-concentrating mechanism increased from 0.19% to 0.33%. Several CCM-associated proteins also increased substantially: PCCAB1 (CA4), PCCAB2 (CA7), and PCCAA1 (CAII) increased 13.7-, 4.1-, and 5.3-fold, respectively, while an LCIB/LCIC-family protein increased 5.1-fold. CA4 became the most abundant carbonic anhydrase in air.

The largest decline in proteome allocation occurred in translation, whose share fell from 16.3% under high CO_2_ to 11.0% in air. This reduction was concentrated in the cytosolic machinery, with allocation to cytosolic ribosomal proteins decreasing by approximately 43%, from 8.3% to 4.8%, and translation factors decreasing from 4.2% to 3.0%. By contrast, plastid ribosomal proteins maintained a nearly constant share, at 1.9% under high CO_2_ and 1.8% in air.

Nitrogen assimilation accounted for a smaller share of the proteome in air, decreasing from 2.45% to 1.64%. This response included the machinery for utilizing urea, the sole nitrogen source supplied in the medium. The urea transporter DUR3 decreased 4.9-fold in abundance, urea carboxylase 8.6-fold, and the two allophanate hydrolases 2.5- and 1.9-fold. Other nitrogen-acquisition pathways also declined, with the nitrate transporter, nitrate reductase, and nitrite reductase becoming 36-, 16-, and 5.3-fold less abundant, respectively, and the ammonium transporter AMT1;3 decreasing 3.6-fold. The urea- and nitrate-utilization genes were organized into two compact genomic clusters on separate chromosomes, each spanning less than 9 kb and containing divergently transcribed gene pairs. In contrast to these pronounced reductions in nitrogen acquisition, the central GS–GOGAT pathway, which channels newly acquired and recycled nitrogen into glutamine and glutamate for cellular biosynthesis, changed comparatively little. The two glutamine synthetases decreased 1.8- and 1.3-fold, while neither ferredoxin-dependent nor NADH-dependent GOGAT changed significantly in abundance.

Pathway-level analysis revealed frequent disagreement between transcript and protein responses to CO_2_ limitation (Figure 2G). At a 5% FDR, fry detected significant responses in 33 of the 44 functional categories in the transcriptome and 39 in the proteome, with 29 categories significant in both. CAMERA, which tests each category against genes outside that category, identified only Calvin cycle regeneration as significant in the transcriptome (adjusted *P* = 0.035) and detected no significant categories in the proteome. Of the 29 categories significant in both layers, 17 changed in opposite directions. Transcripts associated with Calvin cycle regeneration, photorespiration, the carbon-concentrating mechanism, and the light-harvesting antenna decreased, whereas the corresponding proteins increased. Conversely, ribosome biogenesis showed the strongest mean transcript increase (+2.55 log_2_FC) but decreased at the protein level (−0.35 log_2_FC), with nucleotide synthesis and cytosolic ribosomal proteins showing the same pattern. The photosynthetic core and chlorophyll biosynthesis decreased at both levels. Thus, increased protein investment in carbon fixation and reduced investment in cytosolic translation were frequently accompanied by opposing transcript responses.

The divergent proteome and transcriptome responses at the pathway level prompted us to examine how closely these two omic layers correlated at the individual-gene level.

Condition-mean transcript abundance and relative molar protein abundance showed moderate correlations across 5,911 unambiguously paired nuclear genes (Spearman ρ = 0.425 under high CO_2_ and 0.364 in air; Figure 2H; Supplementary Data Set 2), comparable to the ρ = 0.45 reported in *Chlamydomonas reinhardtii* (McWhite et al. 2020). However, transcript and protein log_2_ fold changes under CO_2_ limitation were almost uncorrelated across 5,982 unambiguously paired nuclear genes (Pearson *r* = 0.070, R^2^ = 0.005; Figure 2I; Supplementary Data Set 2). Near-zero correlations between transcript and protein responses have been observed under nutrient depletion in *Escherichia coli* (Glover et al. 2022). Thus, despite a moderate relationship between transcript and protein abundances, transcript responses at the sampled stage provided little information about the extensive proteome remodeling under CO_2_ limitation.

### CO_2_ limitation remodels phosphorylation across photosynthesis and carbon metabolism

Protein abundance alone does not capture regulation through post-translational modifications. To examine this additional layer of the response to CO_2_ limitation, we enriched and quantified phosphopeptides from the same preparations used for global proteomics. This analysis yielded 29,076 phosphosite entries on 4,253 proteins, comprising 23,624 phosphoserine, 4,999 phosphothreonine and 453 phosphotyrosine entries (Figure 3A). After correction for protein abundance, 11,079 entries changed significantly, with 5,994 increasing and 5,085 decreasing in air (Figure 3B). CO_2_ limitation therefore elicited widespread phosphorylation changes alongside extensive remodelling of protein abundance.

**Figure 3.**
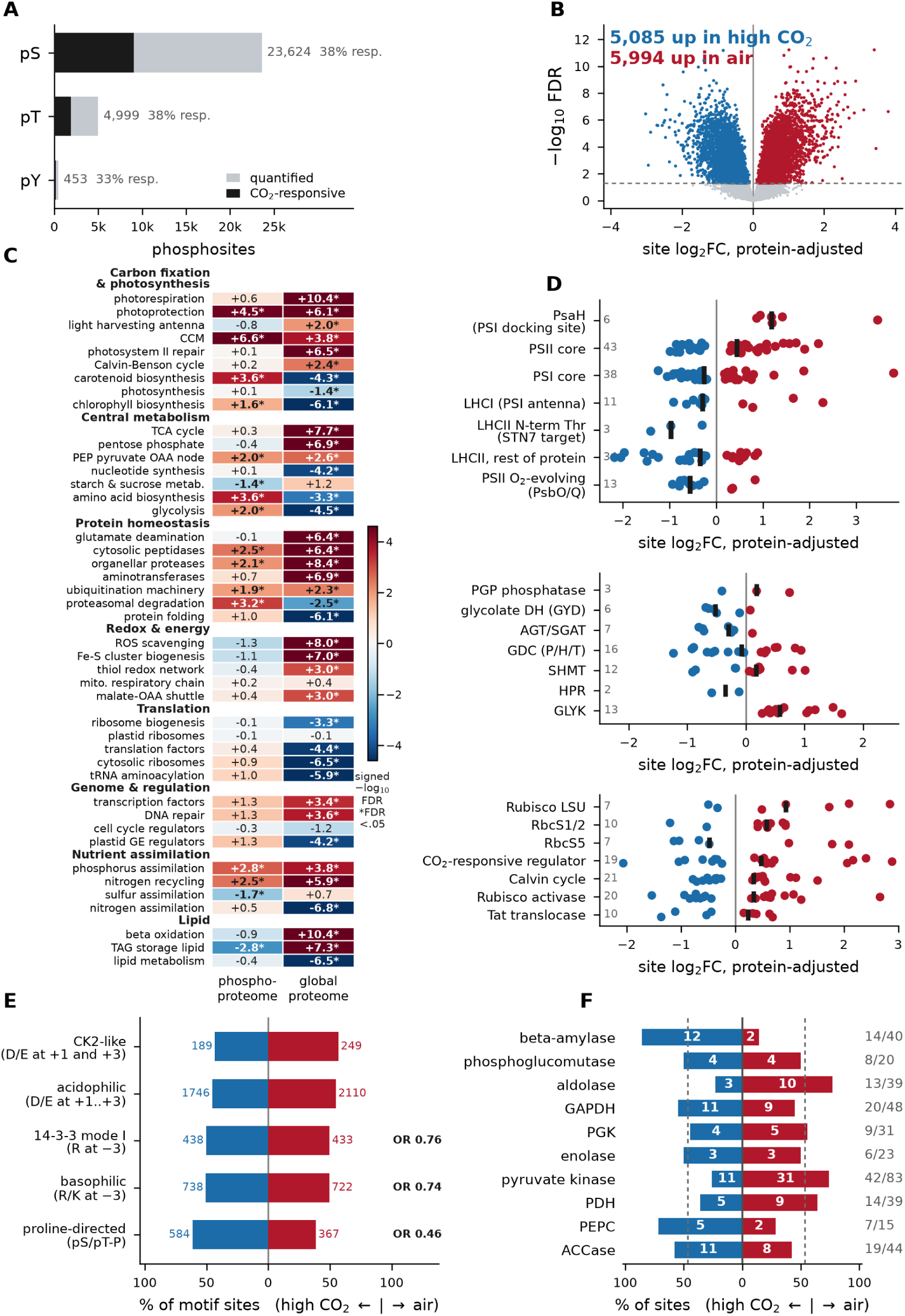
Phosphoproteome response to CO_2_ limitation in *Picochlorum celeri*. Site log_2_ fold changes are protein-adjusted (κ = 0.46) for the air versus high CO_2_ contrast, with red indicating higher phosphorylation in air and blue indicating higher phosphorylation under high CO_2_. A) Composition of the phosphoproteome by modified residue, showing all quantified sites and the subset responsive to CO_2_ limitation at a false discovery rate (FDR) < 0.05. In total, 29,076 sites were quantified on 4,253 proteins. B) Volcano plot of protein-adjusted phosphosite log_2_ fold change versus −log_10_ FDR for all quantified sites. The dashed line marks FDR = 0.05. C) Pathway-level responses across the 44 curated gene sets in the phosphoproteome and total proteome, shown as signed −log_10_ FDR values from fry rotation tests. Positive values indicate higher abundance or phosphorylation in air and asterisks mark FDR < 0.05. D) Protein-adjusted phosphosite changes grouped by photosynthetic complex (top), photorespiratory pathway step (middle), and carbon-fixation proteins including Rubisco and its two small-subunit isoforms (bottom). Black ticks mark row medians. Rows in the top sub-panel are grouped by complex, rows in the middle sub-panel follow the order of the photorespiratory pathway, and rows in the bottom sub-panel place the Rubisco subunits first. Points in the top and bottom sub-panels are sites significant at FDR < 0.05 and the gray numbers count them, whereas the middle sub-panel shows every quantified site of each step and the gray numbers count those. E) Direction of significant phosphosite changes within kinase-recognition motif classes, shown as the percentage higher under high CO_2_ or in air, with counts beside each bar. Odds ratios (OR) for the share of sites higher in air relative to the phosphoproteome-wide background are printed for classes significant by Fisher’s exact test at Benjamini–Hochberg (BH) adjusted q < 0.05, and values below 1 indicate a bias toward high CO_2_. Motif classes overlap. F) Direction of significant phosphosite changes across ten reaction steps from starch to malonyl-CoA. Counts are shown within each bar, gray numbers indicate significant versus quantified sites, and dashed lines mark the phosphoproteome-wide directional background. Bars in E and F are normalized to 100% and indicate direction rather than effect size or number of sites. Sites were quantified from three biological replicates per condition, with BH correction applied across all quantified sites.

Pathway-level analysis showed that phosphorylation responses did not consistently track changes in protein abundance (Figure 3C). Across the 44 curated categories, fry detected significant phosphorylation responses in 16 pathways and protein-abundance responses in 39 at 5% FDR. Fourteen pathways changed significantly in both measurements, with phosphorylation and protein abundance moving in opposite directions in six. The carbon-concentrating mechanism and photoprotection showed the strongest statistical support for increased phosphorylation, accompanied by increased protein abundance. By contrast, phosphorylation increased in carotenoid biosynthesis and proteasomal degradation despite declining protein abundance, whereas TAG storage-lipid metabolism showed decreased phosphorylation despite increased protein abundance. These contrasting responses revealed an additional layer of regulation beyond changes in protein abundance.

To identify regulatory patterns shared across pathways, we examined the sequence contexts of 7,842 responsive phosphosites with unambiguous residue assignments (Figure 3E). Proline-directed phosphorylation was preferentially reduced in air. Only 38.6% of these 951 sites increased, compared with 55.5% of all 7,842 sequence-assigned responsive sites (OR = 0.46, p-adj = 8.6 × 10^−29^). This bias extended across 715 proteins and persisted among proteins with little change in abundance. Proline-directed motifs are recognized by cyclin-dependent kinases, which regulate cell-cycle progression, and MAP kinases, which coordinate responses to environmental signals (Songyang et al. 1996). Their preferential decrease in phosphorylation therefore provides a potential link between CO_2_ limitation and signalling changes associated with growth arrest.

Phosphorylation of the light-harvesting antenna regulates excitation-energy distribution between photosystems through state transitions (Lemeille et al. 2010). Under CO_2_ limitation, antenna proteins and PsaH, a PSI subunit required for efficient energy transfer from the mobile antenna, showed opposing phosphorylation responses (Figure 3D, top; Lunde et al. 2000). Three LHCII N-terminal threonines showed decreased phosphorylation (median log_2_FC = −0.97), as did the multiply phosphorylated N-terminal region of CP29. In *Chlamydomonas*, phosphorylated CP29 associates with PSI under state 2 conditions near PsaH, linking these proteins to antenna redistribution (Kargul et al. 2005). Although sequence alignment did not uniquely resolve the corresponding regulatory residue in *P. celeri*, phosphorylation across the CP29 N-terminal region decreased, whereas all six responsive PsaH entries increased (median log_2_FC = +1.19). Phosphorylation also decreased at ten of 13 responsive entries on the oxygen-evolving subunits PsbO and PsbQ (median log_2_FC = −0.56), whereas the PSII core showed mixed responses. Together, these changes suggest a role for phosphorylation in regulating antenna assembly and function under CO_2_ limitation.

Phosphorylation changes extended to Rubisco and regulators of carbon fixation and growth (Figure 3D, bottom). The Rubisco large subunit showed five significant increases and two decreases. RbcS1/2 showed predominantly increased phosphorylation, with eight increases and two decreases among 12 quantified entries, whereas RbcS5 showed three increases and four decreases among ten entries. Rubisco activase, which promotes Rubisco activation, changed at 20 of 27 quantified entries, although the net direction depended on the strength of the protein-abundance correction. In *Arabidopsis*, phosphorylation of activase Thr78 inhibits its function (Kim et al. 2019). Although this residue is replaced by methionine in *P. celeri*, the adjacent S61 gained phosphorylation in air, identifying a candidate regulatory site near the characterized position.

Phosphorylation also changed at 19 of 33 entries on a putative orthologue of CIA5, a regulator required for induction of the carbon-concentrating mechanism in *Chlamydomonas* (Xiang et al. 2001). A further response involved the CCA1/LHY-like transcription factor, whose TG2 allele increased growth rate by 15% and carbohydrate content by 25% when expressed in TG1 (Steichen et al. 2024). Its protein abundance decreased approximately 1.8-fold in air despite a 1.8-fold increase in transcript abundance, while 15 of 39 phosphosite entries changed significantly, with 12 increasing and three decreasing. These findings extend the phosphorylation response to regulators with established roles in carbon acquisition, growth and carbon storage.

Within photorespiration, phosphorylation increased on enzymes involved in carbon and nitrogen recovery (Figure 3D, middle). Glycerate kinase, which returns photorespiratory carbon to the Calvin cycle, showed 11 significant increases among 13 quantified phosphosite entries. Eight increases occurred within the S203–S214 region, where six of the 12 amino acids were detected as phosphorylated, identifying a densely phosphorylated candidate regulatory region. The glycine decarboxylase complex, which releases CO_2_ and ammonia during photorespiration, showed mixed responses, with seven significant increases and six decreases among 16 entries. Plastid glutamine synthetase, which reassimilates the released ammonia, showed seven significant increases, six within the T312–S336 region. Neither glycerate kinase nor plastid glutamine synthetase showed significant decreases. By contrast, glycolate dehydrogenase and the aminotransferases showed four and three significant decreases, respectively. These contrasting responses suggest selective phosphorylation-dependent regulation of photorespiratory carbon and nitrogen recovery, although the effects of these sites on enzyme activity remain uncharacterized.

Phosphorylation changes also affected enzymes controlling pyruvate production and utilization. Pyruvate phosphate dikinase (PPDK), which can convert pyruvate to phosphoenolpyruvate (PEP), increased 3.8-fold in abundance, whereas all three pyruvate kinases, which catalyze the opposing reaction, declined. PPDK also showed three significant phosphorylation decreases, including at S516. This residue corresponds to maize Ser528, a phosphorylation target of PPDK regulatory protein that also contributes structurally to phosphorylation of the adjacent inhibitory threonine (Chen et al. 2014). Phosphorylation of that threonine, T515 in *P. celeri*, did not change significantly, leaving the effect on PPDK activity unresolved. Mitochondrial pyruvate dehydrogenase E1α gained phosphorylation at S295, corresponding to the conserved inhibitory site and suggesting restricted conversion of pyruvate to acetyl-CoA (Figure 3F; Hirani et al. 2011). Pyruvate kinases, PEP carboxylase and PEP carboxykinase also showed phosphorylation changes, although the functions of their measured sites remain uncharacterized. Together, increased PPDK abundance, reduced pyruvate-kinase abundance and phosphorylation at the inhibitory PDH site suggest a shift toward PEP regeneration that could support gluconeogenesis, although net flux was not measured.

Plastid phosphoglucomutase also gained phosphorylation at S88, corresponding to a regulatory serine whose phosphorylation inhibits the enzyme and restricts glycogen mobilization in *Synechocystis* (Doello et al. 2022). Increased phosphorylation at this conserved position suggests that CO_2_ limitation may engage a regulatory mechanism controlling carbon exchange between starch and central metabolism.

### Ubiquitination as a CO_2_-responsive post-translational modification

Nearly 6% of the *Arabidopsis* genome encodes components of the ubiquitin– proteasome system, including more than 1,400 ligase loci (Vierstra 2009). *P. celeri* devotes 3.6% of its 7,488 nuclear genes to this system and encodes about 150 ligase components, predominantly RING-type enzymes, with only 15 F-box genes compared with approximately 700 in *Arabidopsis* (Gagne et al. 2002; Supplementary Data Set 4). We profiled ubiquitination to examine its association with protein loss in air and transcript–protein uncoupling, and to identify candidate sites of metabolic regulation.

Sites quantified in both conditions were assessed through protein-abundance-corrected fold changes, as in the phosphoproteome, while a binary analysis identified sites lost or gained between conditions. Both analyses showed a collapse in ubiquitination under CO_2_ limitation (Figure 4A). The decline was evident before modeling, with 52–61% of K-GG reporter ions below detection in air compared with 34–37% under high CO_2_, and a 0.65 log_2_ decrease in summed K-GG signal per channel. Of 3,498 quantified sites, 372 decreased and 111 increased. The binary analysis showed a stronger, 15-to-1 asymmetry, with 391 sites lost in air and 26 gained among 681 candidates. A further 163 sites were too close to the threshold to classify. The quantified sites mapped to 1,572 proteins, including 763 with multiple sites and one with 34 (Figure 4B).

**Figure 4.**
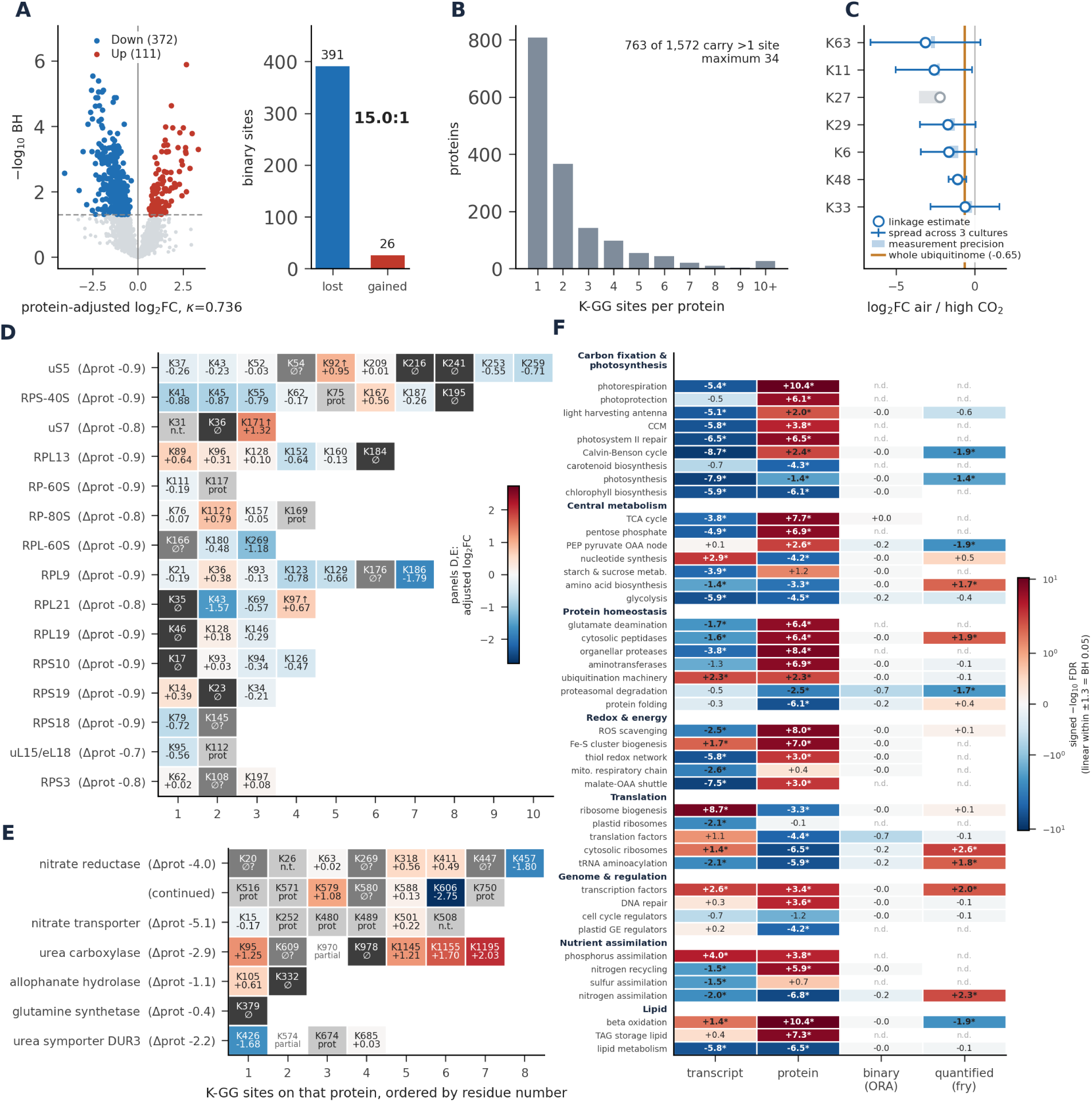
Ubiquitination response to CO_2_ limitation in *Picochlorum celeri*. Wild-type cultures were compared between air and high CO_2_ with three biological replicates per condition. Log_2_ fold changes are air relative to high CO_2_, and quantitative diglycine-remnant (K-GG) site changes were adjusted for the protein-abundance change with κ = 0.736. A) Quantitative and binary components of the ubiquitination response. Left, volcano plot of 3,498 quantified K-GG sites, with colored sites passing Benjamini– Hochberg (BH) adjusted p < 0.05 at κ = 0.736. Of these, 372 were lower and 111 higher in air. Right, sites detected only under one condition, with 391 lost and 26 gained in air. Quantified and lost/gained sites are shown separately because sites detected in only one condition have no fold change. B) Distribution of quantified K-GG sites per protein across the 1,572 proteins with a quantified site. C) Changes in the seven ubiquitin chain linkages estimated from the 661 linkage-diagnostic peptide-spectrum matches (PSMs). Open circles are the per-linkage estimates, whiskers the spread across the three cultures, and pale boxes the precision of the PSM-level estimate. The orange line marks the whole-ubiquitinome shift of −0.65 log_2_ units, and K27 is grayed because it rests on three PSMs. D) Site-level ubiquitination across 15 cytosolic ribosomal proteins containing both quantified and vanished sites. E) Corresponding analysis for six nitrogen-assimilation proteins. In D and E, cells represent individual K-GG sites and row labels give the corresponding protein-abundance change. Vanished sites, detected in one condition only, carry no fold change and are classified by the censoring test: ∅Ø on a dark cell marks a genuine loss, ∅Ø? on a mid-gray cell an indeterminate call, prot on a light-gray cell a loss explained by the protein-abundance change, and n.t. a site that could not be tested. The label partial marks a site detected in only some replicates of a condition, and up-arrows mark sites that also increased in raw intensity. F) Responses of the 44 curated pathways in the transcriptome, proteome, the binary ubiquitination arm (over-representation of lost or gained sites by one-sided Fisher’s exact test, signed by gained minus lost) and the quantified ubiquitination arm (fry). Cells show signed −log_10_ false discovery rate (FDR) with BH adjustment within each column, positive values indicate higher signal in air, asterisks mark FDR < 0.05, and n.d. marks pathways without a result. The color scale is linear within ±1.3 (FDR = 0.05) and logarithmic beyond.

The ten most heavily modified proteins carried 14–34 sites each (Supplementary Data Set 4). The nuclear-pore anchor TPR carried 34, elongation factor 1α 19, and a plasma-membrane H^+^-ATPase, elongation factor 2 and Hsp90 18 each. The ubiquitin-activating enzyme E1 carried 16, a second plasma-membrane H^+^-ATPase 15, and clathrin heavy chain and two uncharacterized proteins 14 each. These proteins act in translation, protein folding, membrane transport and vesicle traffic. Their mean protein mass fractions across conditions ranked from 2nd to 1,062nd, placing heavily ubiquitinated proteins across a broad abundance range. Of their 180 quantified sites, 25 changed under CO_2_ limitation, with 19 decreasing and six increasing. Five proteins carried sites ubiquitinated at corresponding positions in *Arabidopsis* (Song et al. 2024). These comparisons included detected sites outside the quantified set. Hsp90 shared 16 of its 19 conserved lysines and elongation factor 2 shared 12 of 15, while E1, clathrin heavy chain and the H^+^-ATPase shared four, three and two sites, respectively. TPR and elongation factor 1α had *Arabidopsis* orthologues but no shared site, while the remaining three proteins lacked a reciprocal-best-hit orthologue.

Ubiquitination was also detected at 30 lysines on 13 proteins encoded by the chloroplast genome (Supplementary Data Set 4). These included RbcL, the PSI core subunits PsaA and PsaB, the PSII subunits D2 and CP43, four ATP synthase subunits, cytochrome f and the plastid ribosomal protein bL12c.

We next examined whether the ubiquitination response supported increased proteasomal degradation as an explanation for protein loss in air. Proteasome subunits decreased as a set, whereas cytosolic peptidases and organellar proteases increased (Figure 2G). Polyubiquitin protein products, quantified together because they share tryptic peptides, decreased by 0.48 log_2_. K48-linked chains, which are closely associated with proteasomal degradation, also decreased by 1.1 log_2_, together with all six other lysine-linked chain types across 661 spectra (Figure 4C). The largest decrease occurred in K63-linked chains, associated with signaling and membrane trafficking, at 3.1 log_2_, and the smallest in K33-linked chains, at 0.6 log_2_. No linkage differed significantly from the overall ubiquitinome decrease of 0.65 log_2_. Thus, protein loss in air was accompanied by a broad decline in ubiquitination, including degradation-associated K48 chains, and a shift in proteolytic machinery toward peptidases and organellar proteases. This pattern did not support a global increase in proteasomal targeting, although the relative contributions of protein synthesis and degradation to transcript– protein uncoupling could not be resolved from these measurements alone.

Against the global decline in ubiquitination, pathway analysis revealed selective increases in nitrogen assimilation and cytosolic ribosomal proteins (Figure 4F). Across the same 44 curated pathways examined in the other omics layers, 11 of the 23 pathways with sufficient quantified sites showed significant ubiquitination responses. Ubiquitination decreased on Calvin cycle regeneration and the phosphoenolpyruvate– pyruvate–oxaloacetate node, whose proteins became more abundant, and on proteasomal degradation. By contrast, ubiquitination increased on nitrogen assimilation and cytosolic ribosomal proteins, which declined in protein abundance (Figures 4D and 4E), potentially indicating selective targeting for degradation. The direction of ubiquitination change matched protein abundance in four of the 11 responding pathways and transcript abundance in five. In the binary analysis, the 391 genuine site losses were distributed across nearly every pathway, with no significant enrichment.

Translation factors and proteasomal degradation showed the strongest enrichment trends, with 12 and 10 sites lost, respectively, and none gained. Among transcription factors, four sites on the type-B response regulator ARR14 increased by 1.1–1.8 log_2_ while protein abundance remained unchanged, providing a clear example of ubiquitination responding independently of protein abundance (Supplementary Data Sets 2 and 4).

All 15 cytosolic ribosomal proteins carried both disappearing and persisting sites, including nine confirmed ubiquitination losses, while a subset of quantified sites increased (Figure 4D). Nitrogen-assimilation proteins likewise showed increased ubiquitination at five of seven quantifiable sites, together with confirmed losses at urea carboxylase K978, allophanate hydrolase K332 and glutamine synthetase K379 (Figure 4E). Increased ubiquitination at some sites therefore coexisted with selective losses at others within two systems whose reduced abundance accompanied the collapse in biomass production.

Comparison with the *Arabidopsis* ubiquitinome revealed a conserved subset of *P. celeri* ubiquitination sites (Song et al. 2024). Across the full set of 4,740 detected sites, 3,267 occurred on proteins with an *Arabidopsis* orthologue, and 1,408 aligned to a conserved lysine. Among these, 296 were also ubiquitinated in *Arabidopsis*, compared with 169 expected by chance among conserved lysines within the same proteins (Supplementary Data Set 4). The shared sites occurred on 177 proteins involved in protein folding, translation, ubiquitin-dependent degradation, vesicle trafficking and central carbon metabolism. Conservation differed markedly between the cytosolic ribosome and nitrogen assimilation (Figures 4D and 4E). Of the 64 ribosomal sites, 51 (80%) aligned to a conserved lysine and 18 (28%) were also ubiquitinated in *Arabidopsis*, distributed across ten of the 15 proteins. Among the 35 nitrogen-assimilation sites, only eight (23%) aligned to a conserved lysine, and only nitrate reductase K579 was shared. Eleven sites could not be evaluated because urea carboxylase, the two allophanate hydrolases and plastid glutamine synthetase had no assigned *Arabidopsis* orthologues in our mapping. Even nitrate reductase and the nitrate transporter, which had clear orthologues, retained few modified lysines.

Proteins carrying shared sites showed greater decreases in ubiquitination than proteins without them, with median protein-level changes of −0.30 and −0.12 log_2_, respectively. However, within the 112 proteins carrying both shared and unshared sites, the two classes did not differ significantly in their responses. Seven shared positions on six proteins overlapped annotated binding regions or modification sites, providing specific candidates for functional regulation. Ubiquitination of K245 on a pyrophosphate-driven proton pump decreased by 2.08 log_2_ while the protein increased in abundance. This position aligned to an annotated substrate-binding lysine, suggesting a potential connection between ubiquitination and proton transport during acclimation. Shared sites also included RbcL K177 and K201, which participate in substrate binding and catalytic activation. Ubiquitination at RbcL K201 would preclude the carbamylation required for activation, potentially marking an inactive RbcL population. Neither RbcL site changed significantly between conditions, and the effects of ubiquitination on Rubisco and proton-pump activity remain untested.

### Lysine as a versatile post-translational modification receptor

To identify candidate CO_2_-binding lysines, we used LysCarComp–MS, which compares cyanate labeling of paired lysate aliquots in the presence of bicarbonate or sodium chloride (King et al. 2022). Reduced labeling under bicarbonate is consistent with protection of the lysine amine by reversible CO_2_ carbamylation. Enrichment with an anti-acetyl-lysine antibody, which also captures homocitrulline, recovered 2,777 cyanate-labeled lysines on 1,581 proteins, approximately a quarter of the detected proteome (Supplementary Data Set 4). Among the 1,361 sites quantified in both aliquots of at least 12 lysates, plex-adjusted limma analysis identified 86 protected sites and 97 that became more reactive under bicarbonate, relative to the overall homocitrulline-labeling response within each lysate. Twenty protected sites also showed responses more than two standard deviations above the mean site response, defining an additional subset with pronounced protection.

Photosynthesis, lipid metabolism and the pentose phosphate pathway showed significant protection at the pathway level (Supplementary Data Set 3). Among the sites with pronounced protection, isocitrate dehydrogenase K102 and K125 and transaldolase K129 identified candidates in central carbon metabolism, while uroporphyrinogen decarboxylase K279 extended the response to tetrapyrrole biosynthesis. Protection at FtsH2 K143 placed a candidate site on a plastid protease whose homolog participates in PSII repair in *Chlamydomonas reinhardtii* (Malnoë et al. 2014). The protected subset also included ubiquitin C-terminal hydrolase K168 and sites on proteasome subunits, identifying candidate CO_2_-binding positions within the machinery that removes ubiquitin and degrades proteins. Histone H2B K4 and elongation factor 2 K484 extended these candidates to chromatin and translation. RbcL K252 was significantly protected but remained just below the additional effect-size cutoff. Neither the activating lysine RbcL K201 nor the lysines of ubiquitin itself were covered in the competition assay, leaving these established CO_2_-binding positions unevaluated by this approach.

Cytosolic ribosomal proteins showed the opposite pathway response, with increased lysine reactivity under bicarbonate. Ribosomal sites accounted for 27 of the 97 exposed sites (28%), compared with 6.3% of nonsignificant sites (Supplementary Data Set 4).

Exposed sites were associated with higher protein abundance and a higher lysine fraction. Protected sites had significantly fewer acidic residues within three positions of the modified lysine, averaging 0.30 compared with 0.62 around nonsignificant sites, and more basic residues, averaging 0.95 compared with 0.79. Nearby acidic and basic residues can alter lysine pKa and thereby its reactivity toward CO_2_, which reacts with the unprotonated ε-amino group (Kesvatera et al. 1996; Gannon et al. 2024).

Cyanate also forms intracellularly through spontaneous decomposition of carbamoyl phosphate and urea (Hagel et al. 1971; Guilloton and Karst 1987). Plants and cyanobacteria detoxify cyanate through a bicarbonate-dependent cyanase, but no cyanase is annotated in the *P. celeri* genome (Espie et al. 2007; Qian et al. 2011). We therefore examined samples analyzed without cyanate treatment for homocitrulline, the stable product of cyanate modification of lysine. Across the global, phosphopeptide-enriched and ubiquitin-enriched proteomes, homocitrulline accounted for less than 0.5% of spectra and mapped to 123 sites on 99 proteins (Supplementary Data Set 4). RbcL carried eight sites, including K201, whose reversible carbamylation by CO_2_ is required for Rubisco activation (Lorimer 1981). Homocitrullination occupies the same ε-amino group and would preclude this activating modification. CP29, CP26, elongation factor 1α and histone H3 carried four sites each, with additional sites on cytosolic glutamine synthetase and the ammonium transporter. In the unenriched proteome, homocitrulline abundance relative to unmodified peptides from the same proteins was 25% lower in air.

The anti-acetyl-lysine enrichment also revealed widespread acetylation, with 4,654 sites on 1,849 proteins, approximately 30% of the quantified proteome (Supplementary Data Set 2). Acetylation was biased toward abundant proteins across compartments, and 51 of the 99 proteins carrying homocitrulline were also acetylated. The most extensively acetylated proteins carried 14–22 sites each and included enzymes of carbon and nitrogen metabolism, notably RbcL, plastid aldolase, transketolase, phosphoglycerate kinase, ferredoxin-dependent glutamate synthase and carbamoyl-phosphate synthetase. Extensive modification also occurred on proteins involved in translation, folding and trafficking, including elongation factors 1α and 2, DnaK2, ClpC1, protein disulfide isomerase and clathrin heavy chain, alongside one uncharacterized protein.

Despite its breadth, the acetylation response to air was selective, with 113 sites on 107 proteins changing significantly after adjustment for protein abundance (Supplementary Data Set 2). Acetylation increased in nitrogen assimilation and nucleotide synthesis, whereas it decreased in photorespiration, ribosome biogenesis and iron–sulfur cluster assembly. Several of the largest decreases occurred on chromatin-associated proteins. Sites on both subunits of the histone chaperone FACT, a nucleoplasmin-like protein, an inhibitor of growth 1 homolog and a histone H3 lysine-4 methyltransferase showed 3.7- to 7-fold decreases in acetylation without corresponding changes in protein abundance.

RbcL carried 17 acetylated lysines, including K14, K18, K146, K175, K252, K316, K356 and K463, previously identified as acetylated in *Arabidopsis* (Finkemeier et al. 2011).

K334, another reported plant RbcL acetylation site, was also detected (Gao et al. 2016). Only K161 changed significantly under CO_2_ limitation, with acetylation increasing approximately fourfold in air after adjustment for RbcL abundance (Supplementary Data Set 2). K252 was both acetylated and protected in the CO_2_ competition assay, whereas K201, detected as ubiquitinated and homocitrullinated in the preceding analyses, was not detected as acetylated.

The *P. celeri* acetylome contained approximately 3.4 times as many sites as the published *Chlamydomonas reinhardtii* catalogue (Füßl et al. 2022). Comparison between the two species identified 226 sites on 135 proteins that were acetylated at corresponding lysines (Supplementary Data Set 4). Shared sites occurred more frequently than expected among conserved lysines within the same proteins, with a Mantel–Haenszel odds ratio of 3.68, similar in magnitude to the enrichment observed for conserved ubiquitination sites. These positions occurred on proteins involved in carbon metabolism and translation, including plastid aldolase, phosphoglycerate kinase, elongation factors 1α and 2, and ribosomal proteins. Seven shared sites changed significantly in acetylation under CO_2_ limitation.

### The metabolome and lipidome contextualize the post-translationally modified proteome under low CO_2_

To place the proteome and post-translational modification responses in the context of cellular metabolism, we measured the metabolome and lipidome of the same cultures. Gas and liquid chromatography coupled to mass spectrometry quantified 99 metabolite features and 237 lipid species (Supplementary Data Set 2). Forty metabolite features and 104 lipid species differed between air and high CO_2_, with decreases predominating in both layers.

The metabolite response centered on nitrogen metabolism, with depletion extending from arginine precursors to nucleotides and amino sugars (Figure 5B). Ornithine and citrulline were approximately fivefold lower in air, while glutamine decreased by 44%. Complementary analysis of whole-cell hydrolysates from a second cultivation experiment showed that arginine, which was not quantified by untargeted metabolomics, decreased by 52% per cell and from 5.8 to 4.0 mol% of measured amino acids (Supplementary Data Set 1). This was the largest compositional decrease, accompanied by smaller reductions in the glutamate/glutamine and isoleucine fractions. Nucleosides and AMP were three- to sevenfold lower, accompanied by a 2.5-fold decrease in ribose 5-phosphate, whereas aspartate, a precursor for pyrimidine synthesis, accumulated twofold. Glucosamine 6-phosphate and UDP-N-acetylglucosamine, intermediates in glutamine-dependent amino-sugar synthesis, were also depleted, whereas UDP-glucose accumulated threefold. Depletion extended to the cysteine precursor O-acetylserine and the lipid precursors myo-inositol and phosphoethanolamine. Selected carbon-metabolism intermediates accumulated despite these decreases, with sedoheptulose 7-phosphate and fructose 6-phosphate increasing 4.5- and 2.5-fold, respectively.

**Figure 5.**
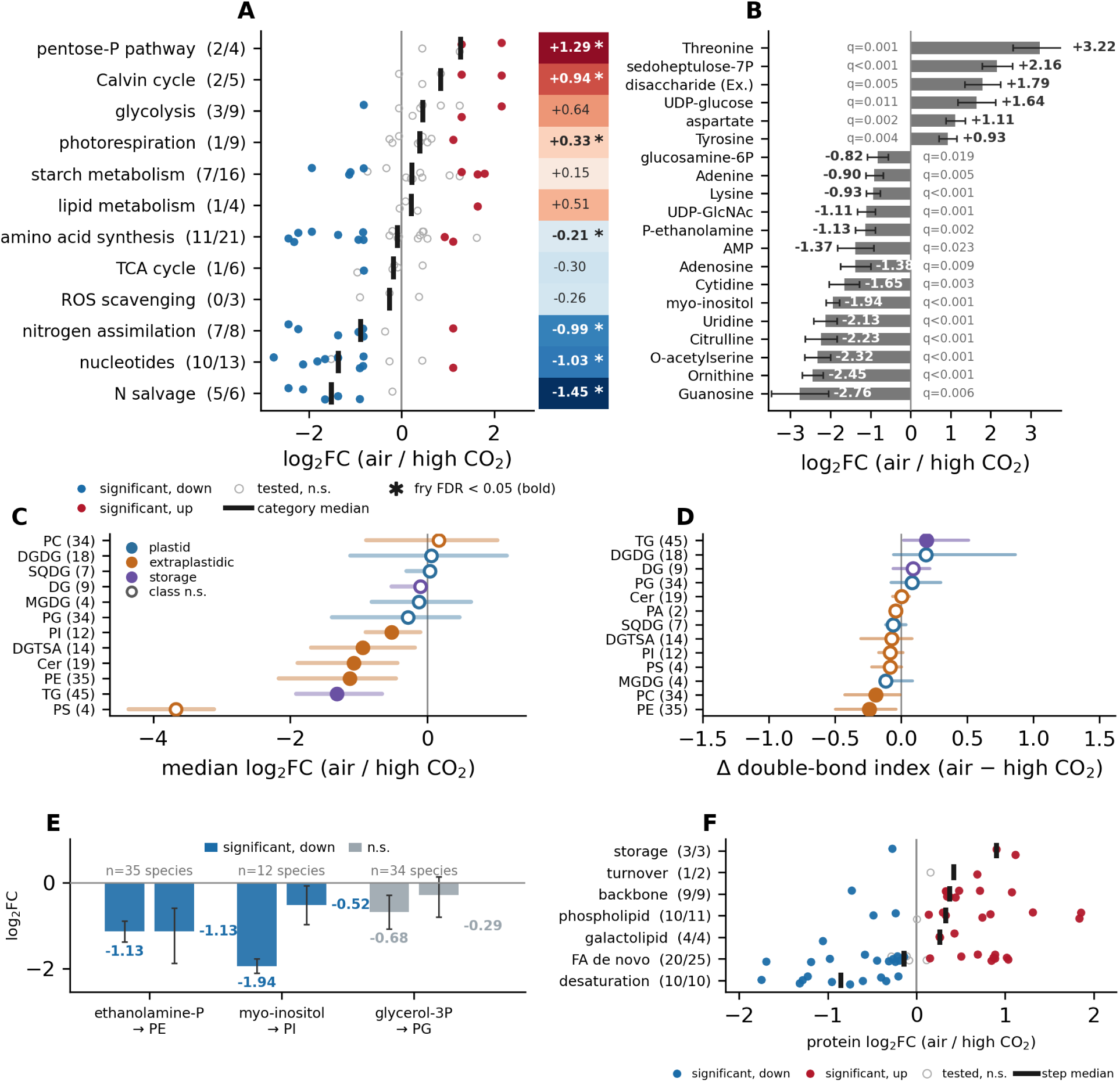
Metabolome, lipidome and lipid-metabolism proteome of Picochlorum celeri under CO_2_ limitation. Three biological replicates were analyzed per condition. Log_2_ fold changes are air relative to high CO_2_, with positive values indicating higher abundance in air. A) Metabolite responses grouped by curated pathway. Points represent individual metabolites, which may occur in more than one pathway, black ticks show pathway medians, and the adjacent heatmap shows the mean log_2_ fold change of each pathway set. Parenthetical values indicate significant versus tested metabolites, and asterisks mark pathway sets with fry false discovery rate (FDR) < 0.05. B) The 20 pathway-assigned metabolites with the smallest adjusted p-values, ordered by log_2_ fold change. Whiskers show the moderated standard error and adjusted p-values are printed alongside. C) Median log_2_ fold change of each lipid class with interquartile range, colored by compartment. Filled symbols indicate a significant class-level change (Wilcoxon signed-rank test across species, p < 0.05) and parenthetical values give the number of lipid species tested. Phosphatidic acid is omitted because only two species were quantified and no class-level test was defined. D) Change in the mean double-bond index of each lipid class between air and high CO_2_, with 95% bootstrap intervals over species. The index is the number of double bonds per acyl chain weighted by each species’ MS signal within its class. Filled symbols mark classes whose interval excludes zero, and colors follow panel C. E) Changes in three headgroup precursor metabolites shown alongside the median response of the lipid class each precursor supplies. Whiskers show metabolite standard errors and bootstrap intervals for lipid-class medians, and numbers above each pair indicate the number of lipid species contributing to the class median. F) Log_2_ fold changes of 64 lipid-metabolism enzymes grouped by pathway step. Black ticks show step medians and parenthetical values indicate enzymes at adjusted p < 0.05 versus quantified enzymes. Lipid classes: Cer, ceramide; DG, diacylglycerol; DGDG, digalactosyldiacylglycerol; DGTSA, diacylglyceryl-trimethylhomoserine; MGDG, monogalactosyldiacylglycerol; PA, phosphatidic acid; PC, phosphatidylcholine; PE, phosphatidylethanolamine; PG, phosphatidylglycerol; PI, phosphatidylinositol; PS, phosphatidylserine; SQDG, sulfoquinovosyldiacylglycerol; TG, triacylglycerol. FA, fatty acid.

At the pathway level, metabolite sets associated with nitrogen assimilation, nitrogen salvage and nucleotide metabolism were approximately 2- to 2.7-fold lower in air (Figure 5A). By contrast, metabolites assigned to the Calvin cycle and pentose phosphate pathway increased, with sedoheptulose 7-phosphate and fructose 6-phosphate contributing to both responses. Metabolites associated with amino acid synthesis showed a modest overall decrease despite opposing responses among individual compounds. Glycolysis, starch metabolism, the tricarboxylic acid cycle and reactive-oxygen scavenging showed no significant changes in their metabolite sets.

Air-grown cells showed pronounced losses of most extraplastidic lipid classes and of triacylglycerol, while none of the four thylakoid lipid classes changed at the class level (Figure 5C). All four phosphatidylserine species were lower in air, by 12.7-fold at the median, and phosphatidylethanolamine was 2.2-fold lower, while the betaine lipid diacylglyceryl-trimethylhomoserine, ceramide and triacylglycerol declined 1.9- to 2.5- fold. Phosphatidylcholine did not change as a class and was retained relative to phosphatidylethanolamine (Mann–Whitney p < 0.001). Within the thylakoid classes, monogalactosyldiacylglycerol (MGDG), digalactosyldiacylglycerol (DGDG) and phosphatidylglycerol contained species that changed in both directions and sulfoquinovosyldiacylglycerol (SQDG) contained none, whereas every changed species of phosphatidylethanolamine, ceramide and the betaine lipid decreased (Supplementary Data Set 2). Across all species, the 63 thylakoid lipids had a median log_2_ fold change of −0.20 against −0.85 for the 174 species of the other classes (Mann–Whitney p = 0.0004).

Phosphatidylcholine and phosphatidylethanolamine became more saturated in air (Figure 5D). Their double-bond indices, the signal-weighted number of double bonds per acyl chain, decreased by 0.20 and 0.24, respectively. Phosphatidylcholine therefore changed in composition despite maintaining its abundance. None of the four major thylakoid lipid classes showed a significant change in double-bond index, whereas triacylglycerol became more unsaturated, gaining 0.19 double bonds per chain. The increased saturation of phosphatidylcholine and phosphatidylethanolamine was accompanied by lower abundance of all ten fatty acid desaturases, with a median decrease of 1.8-fold (Figure 5F). By contrast, enzymes involved in glycerol-backbone assembly and phospholipid and galactolipid headgroup attachment were 1.2- to 1.3-fold more abundant, while triacylglycerol-synthesis enzymes increased 1.9-fold. Enzymes of de novo fatty acid synthesis showed no consistent direction, making desaturation the only enzyme group with a consistent decrease in abundance.

Lower phosphoethanolamine and myo-inositol pools were accompanied by decreases in their corresponding phospholipids (Figure 5E). Phosphoethanolamine decreased 2.2-fold, closely matching the median decrease among phosphatidylethanolamine species, with overlapping uncertainty intervals. Myo-inositol was depleted more strongly than its corresponding lipid class, decreasing 3.8-fold compared with a median decrease of 1.4-fold among phosphatidylinositol species.

## Discussion

Growth of *P. celeri* stopped after CO_2_ supplementation was withdrawn but resumed within hours of its restoration, indicating that the loss of productivity was rapidly reversible. Ekness et al. (2026) likewise observed declining productivity as pH increased, although their cultures continued to grow at pH 8.5. In that study, CO_2_ was supplied to maintain the prescribed pH or replenish inorganic carbon during pH cycling, and growth at pH 8.5 required prior acclimation (Ekness et al. 2026). Our cultures instead experienced sustained withdrawal of supplemental CO_2_, making the two treatments different in carbon supply despite their similarly alkaline pH. The earlier findings therefore show that *P. celeri* can sustain growth at high pH when carbon is replenished, whereas the arrest observed here indicates that ambient-air supply was insufficient to sustain net biomass production under our cultivation conditions. Although the contributions of carbon limitation and alkalinization cannot be separated in our experiment, the rapid return of growth, together with largely preserved PSII photochemical capacity, supports a reversible quiescent state during the interruption of CO_2_ supplementation.

CO_2_ limitation elicited a strong CCM response in *P. celeri* that was shaped substantially by post-transcriptional regulation. CA4 became the dominant carbonic anhydrase in air through a 13.7-fold increase in protein abundance without significant transcript induction. Its induction was accompanied by increased N-terminal phosphorylation, whose effect on enzyme function remains unknown. CA7 and the LCIB/LCIC-family protein were also induced despite decreases in their transcripts, whereas CAII increased at both levels. These contrasting responses show why transcript measurements alone would underestimate the induction of carbon-acquisition machinery. Additional PTM responses included extensive phosphorylation changes on the putative CIA5 orthologue and LCIB/LCIC-family protein, decreased ubiquitination at sites on CA4 and CA7, and decreased acetylation at LCIB/LCIC K409. These modifications identify candidate regulatory sites on proteins involved in the CCM response. Our findings complement the physiological evidence for a rudimentary CCM reported by Ekness et al. (2026), whose growth-derived whole-cell CO_2_ half-saturation estimate was lower than typical Rubisco values. Nevertheless, this substantial CCM response was insufficient to sustain net biomass production under ambient-air supply. Whether the remaining limitation lies in carbon uptake, retention or fixation cannot yet be resolved. Ekness et al. also considered altered Rubisco properties as an untested alternative explanation for the apparent carbon affinity of *P. celeri*. The switch between divergent RbcS isoforms observed here identifies a molecular mechanism through which Rubisco properties could change during acclimation to CO_2_ limitation.

Rubisco remodeling during CO_2_ limitation involved increased abundance of both subunits and Rubisco activase, together with a switch in the predominant small-subunit isoform. RbcS1/2 decreased 5.3-fold while RbcS5 increased 8.1-fold, reversing their relative contributions. Their combined proteome share increased approximately 2.3-fold, complementing the increases in RbcL and Rubisco activase. The two mature RbcS isoforms share only 69.9% amino acid identity, making them substantially more divergent than the two *Chlamydomonas* isoforms, which differ at four residues (Goldschmidt-Clermont and Rahire 1986). Small-subunit identity can influence Rubisco kinetics, as demonstrated with a divergent tobacco isoform (Laterre et al. 2017). In *Arabidopsis*, temperature-dependent shifts in small-subunit composition also accompany changes in CO_2_ affinity, CO_2_/O_2_ specificity and catalytic turnover (Cavanagh et al. 2023). We therefore propose that *P. celeri* responds to CO_2_ limitation by increasing the abundance of its carbon-fixation machinery while favoring a small-subunit isoform that adjusts Rubisco’s catalytic properties to the reduced CO_2_ supply. Whether holoenzymes containing RbcS1/2 or RbcS5 differ in CO_2_ affinity and catalytic turnover remains to be tested, providing a promising direction for future work on the functional significance of this switch. Alongside these changes in abundance and subunit composition, phosphorylation changed extensively on RbcL, both RbcS isoforms and activase. RbcL also carried 17 acetylated lysines, including the catalytic K175 and substrate-binding K334, although only K161 changed significantly, with acetylation increasing approximately fourfold in air. These modifications provide additional potential routes for regulating Rubisco, whose contributions to its assembly and activity during CO_2_ limitation remain unresolved.

Maintenance of PSII photochemical capacity during growth arrest was accompanied by remodeling of the light-harvesting machinery. Greater allocation to antenna proteins, together with PsbS induction, suggests that cells retained light-harvesting capacity while increasing their capacity to dissipate excess excitation as carbon use became constrained. Phosphorylation changes indicate that antenna organization was also subject to regulation. In *Chlamydomonas*, phosphorylation of LHCII and CP29 contributes to antenna redistribution toward PSI during state transitions (Kargul et al. 2005; Lemeille et al. 2010). In *P. celeri*, phosphorylation instead decreased on LHCII N-terminal threonines and the CP29 N-terminal region, while increasing on PsaH, a PSI subunit involved in association with the mobile antenna (Lunde et al. 2000). These contrasting responses suggest altered regulation of antenna–photosystem interactions, although they do not establish the direction of excitation redistribution. The precise regulatory correspondence also remains uncertain because the CP29 alignment did not uniquely resolve the characterized phosphosites. Thus, preservation of PSII capacity was accompanied by changes in antenna abundance, photoprotection and phosphorylation that may help maintain the photosynthetic apparatus during prolonged restriction of carbon fixation.

The transition from rapid growth to maintenance involved selective remodeling of translation at both the protein and PTM levels. Of the 89 quantified cytosolic ribosomal proteins, 88 decreased in abundance despite widespread increases in their transcripts, whereas plastid ribosomal proteins were comparatively maintained. Ubiquitination of the cytosolic ribosome also changed selectively, with increases at some lysines occurring alongside confirmed losses at others on the same proteins. This pattern indicates that the response involved changes in the modification of the remaining ribosomal protein pool as well as reduced protein abundance. Such changes could influence ribosome assembly, function or turnover, although the roles of the individual sites remain unknown. Decreased acetylation of proteins associated with ribosome biogenesis extends this response to the machinery that produces ribosomes. Together, these findings suggest that reduced cytosolic translation during growth arrest involves regulation of both ribosomal protein abundance and modification state. Preservation of plastid ribosomal proteins provides a complementary route for maintaining photosynthetic function, because plastid translation supplies proteins required for PSII repair, including replacement of the D1 reaction-center subunit (Aro et al. 1993; Li et al. 2018). The contrasting responses of the two translation systems are therefore consistent with reduced investment in biomass production while retaining the capacity to maintain the photosynthetic apparatus.

The broad decline in ubiquitination suggests that CO_2_ limitation reshapes regulation of protein function and fate, encompassing signaling, trafficking and turnover. All seven ubiquitin linkage types decreased, including K63 chains associated with signaling and membrane trafficking and K48 chains associated with proteasomal degradation.

Individual proteins nevertheless showed increases in ubiquitination at some lysines and losses at others, indicating selective remodeling. Changes in proteolytic machinery accompanied this response. Proteasome subunits decreased, whereas cytosolic peptidases, organellar proteases and ATG8-conjugation enzymes increased, suggesting a shift in the routes supporting protein turnover. Autophagy is a plausible contributor to the loss of cytosolic ribosomal proteins, given its established role in ribosome turnover (Kraft et al. 2008), although neither autophagic flux nor ribosomal cargo was measured here. Detection of ubiquitination at 30 lysines on 13 chloroplast-encoded proteins also extends the observed modification beyond proteins synthesized in the cytosol. These targets included RbcL, photosystem core proteins, ATP synthase subunits and a plastid ribosomal protein, placing ubiquitination on components of the photosynthetic machinery retained during growth arrest. Ubiquitination of chloroplast-encoded proteins has also been reported in *Arabidopsis* (Sun et al. 2022), although conjugation inside intact chloroplasts remains debated (van Wijk and Adam 2024; Jarvis et al. 2024). Their detection does not establish where conjugation occurs, but it raises questions about both regulatory and degradative roles of ubiquitination in chloroplast protein maintenance. Together, these findings position ubiquitination as a potential link between the functional reorganization of retained proteins and the selective recycling of cellular material during carbon limitation.

Suppression of nitrogen acquisition despite continued urea supply suggests that nitrogen use was adjusted to the loss of carbon-supported growth. The strong decreases in urea transport and catabolism, together with reduced nitrate- and ammonium-acquisition machinery, would limit the capacity to acquire new nitrogen as demand for biomass synthesis declined. By contrast, the comparatively maintained GS– GOGAT pathway and increased abundance of photorespiratory enzymes indicate continued investment in processing nitrogen already within the cell. This distinction also extended to PTMs. Nitrogen-assimilation proteins showed increased ubiquitination and acetylation at the pathway level, together with selective ubiquitination losses on urea carboxylase, allophanate hydrolase and glutamine synthetase. These changes identify potential regulation of nitrogen acquisition and assimilation beyond the decrease in enzyme abundance. Depletion of glutamine, ornithine and citrulline, together with the reduced arginine fraction in whole-cell hydrolysates, further connects this response to altered cellular nitrogen allocation. Reduced external nitrogen acquisition and cytosolic translation therefore appear to be complementary responses that align nitrogen demand with the diminished carbon supply.

Photorespiration provides a route for recycling nitrogen already inside the cell while recovering carbon from Rubisco oxygenation (Figure 6; Eisenhut et al. 2008; Dao et al. 2025). Glutamate:glyoxylate aminotransferase (GGAT) transfers the amino group of glutamate to glyoxylate, producing glycine and 2-oxoglutarate. The glycine decarboxylase complex (GDC) then releases nitrogen from glycine as ammonia, together with CO_2_, and glutamine synthetase (GS) can recapture the released ammonium into glutamine. The glutamate consumed by GGAT can be replenished by glutamate synthase using glutamine and 2-oxoglutarate, or by transferring amino groups from amino acids released during protein turnover onto the 2-oxoglutarate produced by GGAT. Several aminotransferases increased in air, whereas glutamate synthase remained unchanged, providing a potential connection between protein turnover and this redistribution of nitrogen. Increased abundance of enzymes throughout photorespiration also supports greater capacity for carbon recovery, from removal of the inhibitory 2-phosphoglycolate produced by Rubisco oxygenation to regeneration of 3-phosphoglycerate by glycerate kinase. Phosphorylation changed in both directions across GDC and increased on glycerate kinase and plastid GS, identifying potential regulation at the steps that release and reassimilate ammonium and return carbon to the Calvin–Benson cycle.

**Figure 6.**
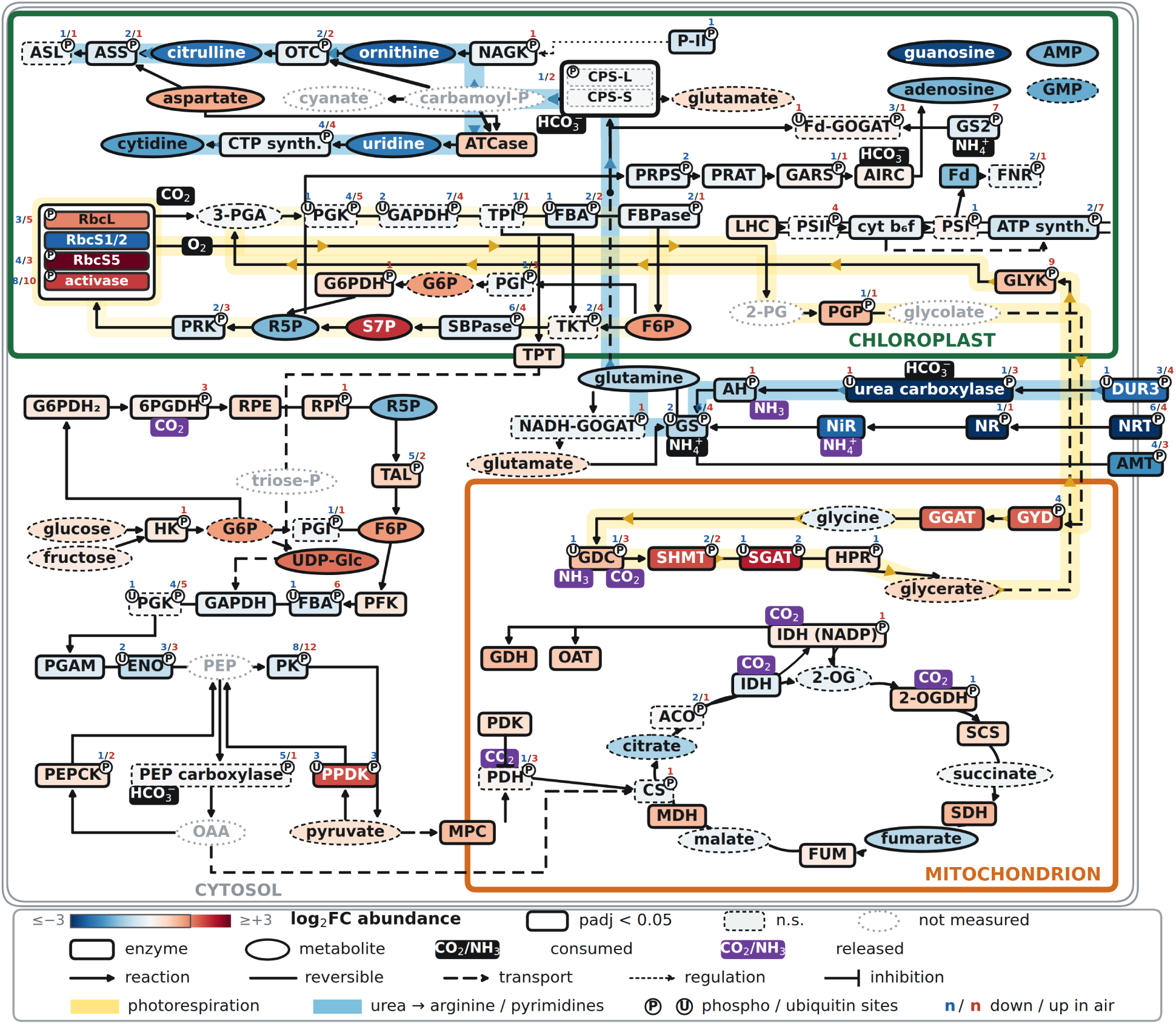
Multi-omic map of carbon and nitrogen metabolism in *Picochlorum celeri* under air relative to high CO_2_. Air is compared with high CO_2_ (air supplemented with 2.25% CO_2_), with positive log_2_ fold change indicating higher abundance in air. Green, orange, and gray borders denote the chloroplast, mitochondrion, and cytosol, respectively. Enzymes are shown as rounded rectangles and metabolites as ellipses. Fill color indicates log_2_ fold change in protein or metabolite abundance, with the scale saturated at ±3. Solid borders indicate adjusted p < 0.05, dashed borders indicate non-significant changes, and gray dotted outlines indicate features not measured. P and U symbols mark phosphosites and ubiquitination sites, respectively, with adjacent blue and red numbers giving the number of residues on the representative protein of each enzyme whose modification decreased or increased significantly in air, each residue counted once in the direction of its largest change (Supplementary Data Set 2). Solid arrows denote reactions, unheaded lines reversible reactions, dashed arrows transport, dotted arrows regulation, and bar-headed lines inhibition. Black labels mark CO_2_, HCO_3_^−^ or NH_4_^+^ consumed by a reaction, and purple labels mark CO_2_, NH_3_ or NH_4_^+^ released. Yellow shading traces photorespiration and its continuation through shared Calvin–Benson cycle regeneration, while blue shading traces urea assimilation through carbamoyl phosphate into arginine and pyrimidine metabolism. Abbreviations: 2-OG, 2-oxoglutarate; 2-OGDH, 2-oxoglutarate dehydrogenase; 2-PG, 2-phosphoglycolate; 3-PGA, 3-phosphoglycerate; 6PGDH, 6-phosphogluconate dehydrogenase; ACO, aconitase; AH, allophanate hydrolase; AIRC, 5-aminoimidazole ribonucleotide carboxylase; AMP, adenosine monophosphate; AMT, ammonium transporter; ASL, argininosuccinate lyase; ASS, argininosuccinate synthetase; ATCase, aspartate carbamoyltransferase; ATP synth., ATP synthase; carbamoyl-P, carbamoyl phosphate; CPS-L/S, carbamoyl phosphate synthetase large/small subunits; CS, citrate synthase; CTP synth., CTP synthase; cyt b_6_f, cytochrome b_6_f complex; DUR3, urea transporter; ENO, enolase; F6P, fructose 6-phosphate; FBA, fructose-bisphosphate aldolase; FBPase, fructose-1,6-bisphosphatase; Fd, ferredoxin; Fd-GOGAT, ferredoxin-dependent glutamate synthase; FNR, ferredoxin-NADP^+^ reductase; FUM, fumarase; G6P, glucose 6-phosphate; G6PDH, glucose-6-phosphate dehydrogenase; G6PDH_2_, cytosolic glucose-6-phosphate dehydrogenase; GAPDH, glyceraldehyde-3-phosphate dehydrogenase; GARS, glycinamide ribonucleotide synthetase; GDC, glycine decarboxylase complex; GDH, glutamate dehydrogenase; GGAT, glutamate:glyoxylate aminotransferase; GLYK, glycerate kinase; GMP, guanosine monophosphate; GS, glutamine synthetase; GS2, plastid glutamine synthetase; GYD, glycolate dehydrogenase; HK, hexokinase; HPR, hydroxypyruvate reductase; IDH, isocitrate dehydrogenase; IDH (NADP), NADP-dependent isocitrate dehydrogenase; LHC, light-harvesting complex; MDH, malate dehydrogenase; MPC, mitochondrial pyruvate carrier; NADH-GOGAT, NADH-dependent glutamate synthase; NAGK, N-acetylglutamate kinase; NiR, nitrite reductase; NR, nitrate reductase; NRT, nitrate transporter; OAA, oxaloacetate; OAT, ornithine aminotransferase; OTC, ornithine transcarbamylase; P-II, nitrogen regulatory protein P-II; PDH, pyruvate dehydrogenase; PDK, pyruvate dehydrogenase kinase; PEP, phosphoenolpyruvate; PEPCK, phosphoenolpyruvate carboxykinase; PFK, phosphofructokinase; PGAM, phosphoglycerate mutase; PGI, phosphoglucose isomerase; PGK, phosphoglycerate kinase; PGP, 2-phosphoglycolate phosphatase; PK, pyruvate kinase; PPDK, pyruvate phosphate dikinase; PRAT, amidophosphoribosyltransferase; PRK, phosphoribulokinase; PRPS, ribose-phosphate pyrophosphokinase; PSI, photosystem I; PSII, photosystem II; R5P, ribose 5-phosphate; RbcL, Rubisco large subunit; RbcS1/2, Rubisco small-subunit isoform encoded by RBCS1 and RBCS2; RbcS5, Rubisco small-subunit isoform encoded by a single gene; RPE, ribulose-phosphate 3-epimerase; RPI, ribose-5-phosphate isomerase; S7P, sedoheptulose 7-phosphate; SBPase, sedoheptulose-1,7-bisphosphatase; SCS, succinyl-CoA synthetase; SDH, succinate dehydrogenase; SGAT, serine:glyoxylate aminotransferase; SHMT, serine hydroxymethyltransferase; TAL, transaldolase; TKT, transketolase; TPI, triose-phosphate isomerase; TPT, triose-phosphate translocator; triose-P, triose phosphate; UDP-Glc, UDP-glucose.

The metabolome and lipidome suggest that growth arrest involved reduced use of carbon for biosynthesis alongside continued expenditure on maintenance. Accumulation of selected Calvin–Benson cycle and pentose phosphate intermediates, together with depletion of nucleotides and amino sugars, is consistent with reduced consumption of central metabolites by growth-associated pathways, without requiring an increase in carbon fixation. Changes in enzyme abundance and phosphorylation suggest that carbon routing was also regulated. Increased PPDK abundance and reduced pyruvate-kinase abundance favor the capacity for phosphoenolpyruvate regeneration, while increased phosphorylation at the conserved inhibitory site of mitochondrial pyruvate dehydrogenase suggests restricted pyruvate oxidation. These responses identify potential control over the balance between carbon retention and respiratory use. The decline in triacylglycerol despite increased abundance of its biosynthetic enzymes is consistent with net consumption of storage reserves during maintenance. Membrane remodeling may complement this redistribution of carbon. Species of the four thylakoid lipid classes were comparatively maintained in air whereas species of the other classes declined. Because these four classes form the bilayer in which the photosystems are embedded, their maintenance is consistent with the retained PSII photochemical capacity and the preserved plastid ribosomal proteins. Retention of phosphatidylcholine alongside depletion of phosphatidylethanolamine could favor bilayer organization, because phosphatidylethanolamine has a greater tendency to form non-bilayer structures (Cullis and de Kruijff 1978). Both phospholipid classes also became more saturated, consistent with the decreased abundance of fatty acid desaturases. A similar combination of increased phosphatidylcholine relative to phosphatidylethanolamine and reduced unsaturation occurs in wheat pollen exposed to high temperature, providing a precedent for coordinated adjustment of membrane composition (Narayanan et al. 2018). Together with the disproportionate loss of non-protein biomass per cell, these observations support a transition in which reduced biosynthetic demand is accompanied by use of stored resources, sparing of the thylakoid lipids and adjustment of the properties of the remaining membranes during growth arrest.

The LysCarComp findings raise the possibility that CO_2_ influences protein function at positions beyond the established catalytic carbamates. Protection of RbcL K252 identifies a candidate interaction within the carbon-fixing enzyme itself, although the absence of K201 from assay coverage prevents comparison with its activating carbamate. Protected sites on ubiquitin-processing and proteasomal proteins extend this possibility to machinery that controls the fate of other proteins. CO_2_-dependent changes in ubiquitin conjugation have been observed in vitro with *Arabidopsis* proteins (Gannon and Cann 2023). CO_2_ could therefore influence cellular processes through interactions with both functional proteins and their regulators. The distinct sequence environments of protected lysines provide a possible chemical basis for this selectivity, because neighboring charged residues can alter lysine pKa and the availability of the unprotonated amine that reacts with CO_2_ (Kesvatera et al. 1996; Gannon et al. 2024). Increased cyanate reactivity requires a different explanation. Bicarbonate binding or carbamylation elsewhere on a protein could change its conformation or local electrostatic environment, making some lysines more accessible or reactive. The assay therefore captures a broader response to bicarbonate than protection alone, and its in vitro measurements cannot establish carbamate occupancy in living cells. These findings nevertheless identify specific sites for testing whether CO_2_-dependent changes in lysine chemistry alter protein activity or interactions, providing a potential route for metabolic conditions to feed back directly on cellular function.

The nitrogen response provides a metabolic context for understanding how cyanate could connect carbon availability to persistent protein modification. Because GDC releases both ammonium and CO_2_, photorespiration could feed a broader recycling loop in which ammonium is recaptured by GS into glutamine while photorespiratory carbon re-enters the bicarbonate pool. Both substrates converge at carbamoyl-phosphate synthetase (CPS), which consumes glutamine, bicarbonate and ATP to produce glutamate and carbamoyl phosphate. The glutamate released by CPS could, in turn, replenish the amino donor used by GGAT, connecting carbamoyl-phosphate synthesis to continued intracellular nitrogen recycling. CPS abundance remained unchanged in air, whereas enzymes consuming carbamoyl phosphate through the arginine branch declined alongside ornithine and citrulline. Arginine also showed the largest decrease in its molar proportion among amino acids measured in whole-cell hydrolysates, connecting suppression of this branch to a change in cellular amino acid composition.

The pyrimidine branch responded differently, with increased aspartate carbamoyltransferase despite depletion of downstream nucleotide pools. These contrasting responses raise the possibility of an imbalance between carbamoyl-phosphate production and consumption. Because carbamoyl phosphate can decompose spontaneously to cyanate, such an imbalance could connect altered nitrogen allocation to cyanate exposure (Guilloton and Karst 1987). Urea provides another potential source of cyanate through spontaneous decomposition (Hagel et al. 1971). Its intracellular availability depends on the balance between import and utilization by urea carboxylase, a reaction requiring bicarbonate and ATP. Both the urea transporter and the enzymes of urea catabolism decreased strongly in abundance in air, identifying another point at which altered nitrogen processing could affect cyanate production. Together, these pathways provide a chemical connection through which the balance between carbon supply, nitrogen processing and energy availability could influence irreversible lysine modification.

We propose that local intracellular CO_2_ availability influences protein function both through reversible carbamylation and through competition with cyanate for susceptible lysine amines. This local CO_2_ environment depends on environmental carbon supply, inorganic-carbon transport, interconversion between CO_2_ and bicarbonate mediated by carbonic anhydrases, and metabolic reactions that produce or consume CO_2_.

Carbamate formation changes the charge and chemical properties of the lysine side chain and can directly regulate protein activity, as demonstrated by the activating carbamate on RbcL K201. Higher local CO_2_ could therefore favor carbamylation, altering protein function while temporarily protecting the same lysines from homocitrullination. The outcome would also depend on cyanate exposure and the protonation state of each lysine, linking modification to both cellular carbon status and the local protein environment. Because carbamylation is reversible, its occupancy can track the current local CO_2_ environment. Homocitrullination, by contrast, is irreversible and can persist for the lifetime of the modified protein (Gorisse et al. 2016), allowing its abundance to integrate exposure to cyanate while the lysine remains unprotected.

Imbalances between urea import and utilization, or between carbamoyl-phosphate production and consumption, could increase that exposure. Susceptible lysines could therefore integrate environmental carbon supply, intracellular carbon and nitrogen metabolism, and protein lifetime through two competing modifications that alter protein function on different timescales.

The protected and exposed lysines identified by LysCarComp provide targets for testing both the functional consequences of modification and their dependence on cellular metabolic state. GDC is particularly informative because it produces CO_2_ and contains sites with contrasting responses, including protected T-protein K413 and exposed L-protein K442 and P-protein K1003. Determining whether modification at these positions affects glycine-cleavage activity, complex assembly or interactions would establish their functional significance and potential roles in regulating a key photorespiratory step.

RbcL provides a complementary set of targets, with protected K252, exposed K183 and the activating K201, at which homocitrullination was detected. Cyanate is known to irreversibly inactivate Rubisco through modification of lysines in RbcL (Chollet and Anderson 1978). Detailed quantification of carbamylated, homocitrullinated and unmodified forms of these sites in vivo across varying CO_2_ concentrations and different metabolic states, including carbon limitation and recovery, would test whether the two modifications change reciprocally as predicted by competition for the same amine.

Relating these measurements to local CO_2_ availability, cyanate exposure and protein renewal would distinguish the chemical and temporal contributions to modification.

Extensive phosphorylation remodeling and selective changes in ubiquitination and lysine acetylation show that the air response extended beyond changes in protein abundance to multiple forms of post-translational regulation. Lysine emerges from these data as a major regulatory hub capable of integrating chemically distinct inputs through both enzymatic and non-enzymatic modification. Its side chain can carry ubiquitin through enzyme-catalyzed conjugation, respond to acetyl-CoA through acetylation, and react directly with CO_2_ or cyanate to form carbamates or homocitrulline. The same residue class can therefore register changes in regulated protein turnover, metabolic state and local chemical environment through modifications with very different mechanisms and lifetimes.

This study presents an extensive multiomics dissection of *P. celeri* under high and low CO_2_, revealing how its physiology changes as rapid growth gives way to reversible growth arrest. By integrating protein abundance and post-translational modifications with gene expression, metabolite pools and cellular phenotypes, we connect the observed loss of productivity to changes across multiple regulatory layers. The resulting dataset lays the groundwork for generating testable mechanistic hypotheses about the processes underlying *P. celeri*’s fast growth and their broader relevance to other photosynthetic organisms.

## Supporting information

Supplementary Data Set 1

Supplementary Data Set 2

Supplementary Data Set 3

Supplementary Data Set 4

## Acknowledgments

This research was performed on project awards from the Environmental Molecular Sciences Laboratory (EMSL) (https://doi.org/10.46936/expl.proj.2024.61507/60012925 and https://doi.org/10.46936/camc.proj.2026.62132/60015765), a DOE Office of Science User Facility sponsored by the Biological and Environmental Research program under Contract No. DE-AC05-76RL01830. We thank Sarah Leichty and Rick Washburn, project managers in the EMSL User Program Services team, for managing the projects that produced this research, and Elisabeth Citta and Cory Smith of the Environmental Health and Safety office of the Colorado School of Mines for procuring reagents and for the safe shipment of samples to EMSL. We also thank the U.S. Department of Energy for its sustained support of algal research, and the managerial and administrative staff at EMSL and at the Colorado School of Mines, whose work behind the scenes made this study possible.

## Use of large language models

Bioinformatic analyses and the editing of the text were facilitated by the large language models Claude Fable 5.1 (Anthropic) and GPT-6 Astra (OpenAI). The authors reviewed and verified every analysis and every sentence and take full responsibility for the content.

