## Supplementary File 1 for "Multiomics dissection of the CO_2_-dependent fast growth of *Picochlorum celeri*": README.txt

Supplementary File 1. Complete model inputs, reference sequences, annotations and original PAM measurements.

The model_inputs folder contains the ten complete sample-level matrices used by the reported models, including all 18 samples or paired lysates. The CA-mutant samples contributed to fitting, although the paper reports the wild-type comparison. RNA_counts_18.tsv contains the integer counts required to reproduce DESeq2. The remaining matrices contain log2 abundances or centered log2 competition ratios, as defined in input_index.tsv. Gene name columns are descriptive labels and must not be treated as measurements. Empty cells denote missing values. sample_map.tsv connects model sample identifiers to the sample and channel table in Data Set 2.

The protein search database and target FASTA are exact analysis inputs. The genome FASTA, alignment GFF3, counting GFF3 and allele maps specify the RNA-seq reference. Gene DNA sequences are extracted genomic intervals including introns, not spliced coding sequences.

Gene descriptions use the local annotation_v11.tsv snapshot, with the previous-name suffix removed in result tables. Displayed gene names additionally use display_names.tsv and curated_gene_names.tsv. The core result builder preserves existing PCCAA/PCCAB carbonic-anhydrase labels. published_gene_annotations.tsv records the labels actually displayed by each result sheet. Annotation source codes identify inherited assignments and do not imply experimental validation in P. celeri.

The PAM export is the complete instrument file. Data Set 1 identifies the selected curves and reports their PSII measurements. Instrument clock times on 26 June are 2 h 35 min ahead of wall-clock time.

SHA256 checksums and the purpose of every file are in file_manifest.tsv.
