## Supplementary Information for "Multiomics dissection of the CO_2_-dependent fast growth of *Picochlorum celeri*"

### 1. Marine Dense medium

The final medium contains per liter approximately 40 g Instant Ocean sea salt, 436 mg urea, 89 mg $KH_2PO_4$, 15 mg $FeSO_4·7H_2O$ and 20.25 mg disodium EDTA, together with trace metals (313.8 µg $MnSO_4·H_2O$, 24.23 µg $CoCl_2·6H_2O$, 48.82 µg $ZnSO_4·7H_2O$, 2.00 µg $CuSO_4·5H_2O$ and 6.81 µg $Na_2MoO_4·2H_2O$) and vitamins (1.54 mg thiamine HCl, 4.4 µg biotin and 4.4 µg cyanocobalamin). Approximately 40 g of Instant Ocean sea salt was dissolved per liter of final medium with 7.5 mL of iron stock, 1 mL of trace-metal stock, 1 mL of phosphate stock, 1 mL of vitamin stock and 2 mL of urea stock, and the medium was filter-sterilized. One liter of vitamin stock contained 1.54 g thiamine HCl and 1 mL each of 4.4 g $L^{-1}$ biotin and cyanocobalamin primary stocks. Related dense-medium formulations are described by Weissman et al. (2018) and Cano et al. (2024).

| Stock | Stock concentration | Addition per liter | Final concentration |
|---|---|---|---|
| $FeSO_4·7H_2O$ | 2 g $L^{-1}$ | 7.5 mL iron stock | 15 mg $L^{-1}$ |
| Disodium EDTA | 2.7 g $L^{-1}$ in iron stock | 7.5 mL iron stock | 20.25 mg $L^{-1}$ |
| $MnSO_4·H_2O$ | 313.8 mg $L^{-1}$ | 1 mL trace stock | 313.8 µg $L^{-1}$ |
| $CoCl_2·6H_2O$ | 24.23 mg $L^{-1}$ | 1 mL trace stock | 24.23 µg $L^{-1}$ |
| $ZnSO_4·7H_2O$ | 48.82 mg $L^{-1}$ | 1 mL trace stock | 48.82 µg $L^{-1}$ |
| $CuSO_4·5H_2O$ | 2.00 mg $L^{-1}$ | 1 mL trace stock | 2.00 µg $L^{-1}$ |
| $Na_2MoO_4·2H_2O$ | 6.81 mg $L^{-1}$ | 1 mL trace stock | 6.81 µg $L^{-1}$ |
| $KH_2PO_4$ | 89 g $L^{-1}$ | 1 mL phosphate stock | 89 mg $L^{-1}$ |
| Urea | 218 g $L^{-1}$ | 2 mL urea stock | 436 mg $L^{-1}$ |
| Thiamine HCl | 1.54 g $L^{-1}$ | 1 mL vitamin stock | 1.54 mg $L^{-1}$ |
| Biotin | 4.4 mg $L^{-1}$ in vitamin stock | 1 mL vitamin stock | 4.4 µg $L^{-1}$ |
| Cyanocobalamin | 4.4 mg $L^{-1}$ in vitamin stock | 1 mL vitamin stock | 4.4 µg $L^{-1}$ |

### 2. Medium pH, total alkalinity and the $CO_2$ content of the gas streams

Cell-free Marine Dense medium was sparged in the vessels under each gas stream for 24 h, the interval between daily dilutions, and then sampled, twice and independently, on the first medium batch (four vessels, three titrations each) and on the second (three vessels, three titrations each) some months apart. The pH was read immediately after harvest with the vessels held at 33 °C, and each total-alkalinity titration then took about 10 min without temperature control, which leaves the alkalinity unaffected because the acid equivalence volume depends neither on temperature nor on $CO_2$ exchange during the titration. Total alkalinity was titrated with 0.5 N sulfuric acid on 100 mL samples to the inflection near pH 4.5 (Ekness et al. 2026), so that alkalinity in meq $L^{-1}$ equals the acid volume in µL divided by 200. The medium equilibrated under unsupplemented air formed a precipitate, whereas the medium under air with 2.25% $CO_2$ remained clear. Precipitation of this medium during storage at room temperature, attributed to iron and reduced by added EDTA, has been documented for the ALGiSIM reservoirs, in which the medium is allowed to precipitate in the tank (Karns 2024). Batch 1 gave 3.36 ± 0.14 meq $L^{-1}$ at pH 7.05 ± 0.10 under 2.25% $CO_2$ and 2.19 ± 0.13 meq $L^{-1}$ at pH 8.23 ± 0.01 under air (12 titrations on four vessels each), and batch 2 gave 3.39 ± 0.14 meq $L^{-1}$ at pH 7.00 ± 0.06 under $CO_2$ and 2.13 ± 0.10 meq $L^{-1}$ at pH 8.21 ± 0.01 under air (nine titrations on three vessels each), so the batches differ by 0.03 meq $L^{-1}$ under $CO_2$ and 0.06 meq $L^{-1}$ under air, within one SD. Every titration, with its pH, alkalinity and derived $CO_2$ values, is listed in Supplementary Data Set 1.

The $CO_2$ mole fraction of each gas stream was back-calculated from the pH and alkalinity of the sparged medium with the carbonate treatment of Ekness et al. (2026). Their Table 1, measured in this medium at 33 °C and an alkalinity of 2.85 meq $L^{-1}$, fixes two conventions, dissolved $CO_2$ = K_H × $pCO_2$ with K_H = 0.0223 mol $L^{-1}$ $atm^{-1}$ and $pCO_2$ = P × x_CO2 with P = 0.820 atm, both read as the median ratios of the table's columns. Apparent constants of the medium at 33 °C, $pK_1$ = 5.92 and $pK_2$ = 9.01, were fitted by least squares to the 14 rows of that table through TA = $[HCO_3^-] + 2[CO_3^{2-}] + [OH^-] - [H^+]$, with $[HCO_3^-] = K_1\, dCO_2/[H^+]$ and $[CO_3^{2-}] = K_2[HCO_3^-]/[H^+]$, which reproduces the table's alkalinity within 10% at every row (root mean square 4%). For each titration the same equations were inverted at the measured alkalinity and pH to give dissolved $CO_2$, then $pCO_2 = dCO_2$/K_H and x_CO2 = $pCO_2$/P, and the two batches were summarized separately. The air stream carried 448 ± 20 ppm $CO_2$ at the time of batch 1 and 451 ± 22 ppm at the time of batch 2, whereas the medium sparged for 24 h with the nominal 2.25% stream held 252 ± 58 µM (batch 1) and 278 ± 42 µM (batch 2) dissolved $CO_2$, which corresponds to 1.38 ± 0.32% and 1.52 ± 0.23% $CO_2$. The SD over titrations carries the pH and alkalinity scatter, whereas the fitted constants add an uncertainty of about 5 to 10% that is common to all samples. Assumptions are that the pH probe reads on the same scale as the probe of Ekness et al. (2026), that the phosphate and borate contributions to alkalinity are absorbed into the empirical constants, and that the K_w term (pK_w taken as 13.7) is below 0.2% of the alkalinity at pH 8.2. Full equilibrium with 2.25% $CO_2$ at 0.82 atm and an alkalinity of 3.36 meq $L^{-1}$ would hold 411 µM dissolved $CO_2$ at pH 6.83, but the cultures were diluted with fresh medium every 24 h, so the 24-h value of 250 to 280 µM (1.4 to 1.5% $CO_2$ equivalent) is the carbon status the cultures experienced, and a medium already near equilibrium with laboratory air is unaffected by this consideration.

#### 3. Photobioreactor operation

The ALGiSIM system is described in full by Karns (2024). The culture vessels are Corning 1396 square 500-mL Pyrex media bottles (Corning, Corning, NY, USA), which expose an illuminated face of about 6.5 by 8.5 cm at 400 mL (Karns 2024). Each bottle stands on a load cell above the thermoelectric module and stir plate and is illuminated on one face by a liquid-cooled panel of cool-white LEDs 14 cm from the glass, with the other three sides covered by an insulating black jacket. Irradiance was calibrated before each run in a culture bottle holding 400 mL of water, with the spherical quantum sensor (US-SQS/L, Heinz Walz GmbH, Effeltrich, Germany) placed at the center of the bottle between the 200 and 300 mL marks and read on a LI-COR quantum meter. Mass flow controllers (Aalborg Instruments and Controls) blended $CO_2$ into compressed house air from the building supply (DAC) in a common gas mixer, and the filtered mixture was distributed to the four vessels through rotameters and entered each culture through a sintered-glass sparger (10 to 16 µm pores, VICI Jour). Sterile medium and deionized water were delivered by peristaltic pumps (Watson-Marlow) from reservoirs filled through 0.2 and 0.1 µm filters, and the culture volume was tracked by the load cell beneath each bottle and converted to volume by the medium density, so that evaporative losses were replaced every 2 h and each daily dilution removed and replaced a set volume. Bottles were autoclaved and the instrument plumbing was sterilized in place with 3% bleach before each run.

#### 4. Flow cytometry acquisition, gating and cell size

Samples of the second run were taken from the four vessels at simulated solar noon on Days 5, 6, 7, 9, 11, 12 and 13 (06:00 to 08:30 wall-clock time, the light script being offset from the wall clock) over the high-$CO_2$, air and recovery phases, diluted 1,000-fold in deionized water and analyzed immediately on the Attune NxT at 200 µL $min^{-1}$ with 200 µL analyzed per sample, forward scatter collected on the photodiode at 360 V, side scatter on the 488/10 nm detector at 300 V and chlorophyll autofluorescence on BL3 (695/40 nm) at 340 V, with an FSC-H threshold of 23,000 and an area scaling factor of 1.04. Event rates were 457 to 969 events $s^{-1}$ with 0.22 to 0.75% coincidence, and 28 files were saved as FCS 3.1. Files were read with FlowKit 1.3.2 (White et al. 2021). The cell gate retained events with FSC-A above 20,000 and SSC-A above 2,000 and no scatter channel at or above the 1,040,000 saturation limit. Per-sample medians of FSC-A, SSC-A and BL3-A were computed after a further pulse-shape gate retaining events with $\text{FSC-A} - 1.0259 \times \text{FSC-H} \leq 4 \times 5{,}551$, using the fitted slope and robust residual standard deviation. This operational gate was not calibrated to distinguish individual cells from doublets. Low-chlorophyll events were pulse-shape gated events with BL3-A below 1,000, a fixed gate that lies under the dimmer lobe of the main population on the air days (median BL3-A about 2,400 on day 9) and above the dim cluster that appears on day 11 (median about 400), so that neither cluster straddles it.

Polystyrene size standards (Flow Cytometry Size Calibration Kit, F13838, Invitrogen, Carlsbad, CA, USA; nominal 1, 2, 4, 6, 10 and 15 µm) were acquired at the same detector voltages and flow rate at a matched event rate, and their FSC-A was multiplied by 1.04/1.07 to correct for the different area scaling factor of the bead acquisition. The 10 and 15 µm beads saturate FSC-A at this gain and the 1 µm bead sits at the acquisition threshold, so the 2, 4 and 6 µm beads were the anchors, with modes at

74,086, 309,470 and 703,973 FSC-A units located by Gaussian kernel density estimation on $\log_{10}$ FSC-A. Over 2 to 6 µm, FSC-A = 10^4.2533 × d^2.051 ($R^2$ = 0.99998), that is forward scatter of polystyrene scales with cross-sectional area, and two further bead acquisitions at other event rates and flow rates bracket the acquisition-dependent systematic at −4.0% and +6.2% in diameter.

Angular scattering of homogeneous spheres was computed with an implementation of the BHMIE algorithm (Bohren and Huffman 1983) validated against the Rayleigh limit, the optical theorem and the Wiscombe reference case, with a wavelength of 488 nm, a medium refractive index of 1.337 and a polystyrene index of 1.604. Because the instrument does not publish its scatter collection angles, they were fitted from the beads following Welsh et al. (2019), Ackleson and Spinrad (1988) and Green et al. (2003). The FSC signal was modeled as the unpolarized scattered intensity integrated over an annulus around the beam with a single gain, and the best window was 1.0 to 28.0° (root mean square residual 0.0009 $\log_{10}$ units), with 13 windows reproducing the beads within 0.01 $\log_{10}$ units and giving diameters within 5% of the primary.

Cell diameter was obtained by interpolating each sample's median FSC-A against the modeled forward-scatter curve for a homogeneous sphere at a fixed real refractive index of 1.41 and an imaginary refractive index of 0.003. The diameter grid spanned 1.6 to 5.0 µm in 0.02 µm steps, with interpolation performed on the monotonically increasing $\log_{10}$ FSC curve. The assumed real refractive index is within the range reported for phytoplankton (Aas 1996), although the homogeneous-sphere approximation limits the interpretation of absolute cell size (Green et al. 2003). Cell volume was calculated as $\pi d^3/6$, and daily diameters are the mean and standard deviation across the retained vessels. The B4 flow-cytometry sample acquired on 30 June was excluded because it had been refrigerated for two days before acquisition. Per-sample and daily values are given in Supplementary Data Set 1 and the calibration in Supplementary Figure 1.

#### 5. Amino acid analysis

Aliquots of $2 \times 10^8$ counted cells were hydrolyzed in 200 µL of 6 N HCl with 1% phenol at 110 °C for 24 h, dried and taken up in norleucine dilution buffer (4 mL for the $CO_2$ samples and 1 mL for the air samples), and 50 µL of each, carrying 2.0 nmol norleucine as internal standard, was injected on the Hitachi LA8080 analyzer with post-column ninhydrin detection. Protein per cell was the summed residue mass of the quantified amino acids. For amounts per gram of cell dry weight, ash was assumed to be 15% of dry weight, close to the 11.2% reported for a *Picochlorum* isolate (Dahmen et al. 2014).

### 6. Proteome extraction, digestion and fractionation

The MPLEx extraction used ice-cold chloroform:methanol:water (8:4:3), Tissue-Tearor homogenization (BioSpec Products, Bartlesville, OK, USA) and centrifugation at 12,000 g for 5 min at 4 °C. The remaining lyophilized material was powdered with 2.8-mm ceramic beads in a Bullet Blender (Next Advance; 30 s at speed 5.65) and lysed in 1 mL of 10% SDS, 50 mM triethylammonium bicarbonate (TEAB), 10 mM NaF, 1% each of phosphatase inhibitor cocktails 2 and 3 (Sigma), 1 mM PMSF, 2 µg $mL^{-1}$ aprotinin, 10 µg $mL^{-1}$ leupeptin, 10 mM sodium butyrate, 2 µM SAHA, 10 mM nicotinamide, 50 µM PR-619 and 0.2 mM iodoacetamide. Protein was reduced with 5 mM DTT at 60 °C for 30 min and alkylated with 10 mM iodoacetamide at 45 °C for 45 min in the dark, phosphoric acid was added to 2.7%, followed by seven volumes of cold 90% methanol in 100 mM TEAB, and the protein was captured on S-Trap Midi columns (ProtiFi, Fairport, NY, USA; C002-MIDIX-0020K) and digested overnight at 37 °C with trypsin (Promega, Madison, WI, USA; V5111) and Lys-C (Santa Cruz Biotechnology, Dallas, TX, USA; SC-360250D), each at 1:50 (w/w). Peptides were desalted on Sep-Pak tC18 cartridges (Waters, Milford, MA, USA).

TMTpro-labeled peptides were fractionated on an Agilent 1200 HPLC (Agilent Technologies, Santa Clara, CA, USA) with an XBridge BEH C18 column (250 × 4.6 mm, 3.5 µm, Waters) at 1 mL $min^{-1}$. Solvent A was 4.5 mM ammonium formate, pH 10, in 2% acetonitrile, and B was the same buffer in 90% acetonitrile, with B rising to 16% over 6 min, to 40% over the next 60 min, to 44% over 4 min and to 60% over 5 min before a 14-min hold, and the 96 fractions were concatenated into 12 by combining every twelfth fraction. Five percent of each fraction was reserved for the global proteome, and the remainder was reconstituted in 200 µL of 80% acetonitrile with 0.1% trifluoroacetic acid, loaded onto Fe(III)-NTA cartridges on an AssayMap Bravo at 6 µL $min^{-1}$, washed with the same solvent and eluted with 1% ammonium hydroxide at 5 µL $min^{-1}$ into formic acid and DDM, and the dried phosphopeptides were reconstituted in 12 µL of 3% acetonitrile with 0.1% formic acid.

### 7. LC-MS/MS acquisition for proteomics

Global, K-ε-GG and acetyl-lysine samples were separated on a nanoAcquity UPLC (Waters, Milford, MA, USA) with an in-house packed column (75 µm inner diameter × 25 cm, 1.7 µm BEH C18 particles, Waters) at 200 nL $min^{-1}$, with 0.1% formic acid in water as mobile phase A and 0.1% formic acid in acetonitrile as mobile phase B. The global fractions were eluted with a gradient from 5% to 19% B over 120 min, whereas the K-ε-GG and acetyl-lysine samples were acquired over 150 min. The column was coupled to an Orbitrap Eclipse Tribrid mass spectrometer (Thermo Fisher Scientific) through a FAIMS Pro interface that cycled through compensation voltages of −45, −60 and −75 V. MS1 scans covered m/z 400 to 1800 at 120,000 resolution with a 50-ms maximum injection time. MS2 spectra were acquired in a cycle-time data-dependent mode at 50,000 resolution with 0.7 m/z quadrupole isolation and HCD at a normalized collision energy of 32, with a first mass of m/z 120, exclusion of singly charged precursors and 30-s dynamic exclusion. Maximum MS2 injection times were 86 ms for the global fractions and 125 ms for the K-ε-GG and acetyl-lysine samples.

Phosphopeptides were separated on a Vanquish Neo UHPLC (Thermo Fisher Scientific) with a PepMap Neo trap cartridge and an EASY-Spray PepMap RSLC C18 column (ES906, 150 µm inner diameter × 15 cm, 2 µm particles, 100 Å pores, Thermo Fisher Scientific) coupled to an Orbitrap Exploris 480. MS1 scans covered m/z 300 to 1800 at 60,000 resolution, and the 20 most abundant precursors of each cycle were fragmented by HCD at a normalized collision energy of 30 with 0.7 m/z isolation and recorded over m/z 110 to 1010 at 30,000 resolution, with singly charged precursors excluded and 45-s dynamic exclusion.

Database searches. Spectra were searched with FragPipe 24.0 (MSFragger 4.4, Kong et al. 2017; Philosopher 5.1.3, da Veiga Leprevost et al. 2020) against the non-redundant P. celeri proteome with reversed decoys and common contaminants, with strict trypsin specificity, up to two missed cleavages and peptide lengths of 7 to 50 residues. The contaminant set was the common-contaminant list added by FragPipe without human ubiquitin (UniProt P62979) and human NEDD8 (Q15843), because both share identical tryptic peptides with their P. celeri counterparts, including the diglycine-modified ubiquitin peptides at K6, K33 and K48, which would otherwise have been assigned to the human proteins. The global proteome search used a precursor tolerance of 20 ppm and a fragment tolerance calibrated to 5 ppm, carbamidomethyl cysteine and TMTpro on lysine as fixed modifications, and methionine oxidation (up to three), protein N-terminal acetylation and TMTpro on peptide N-termini as variable modifications with at most three per peptide. Peptide-spectrum matches were rescored with Percolator (Käll et al. 2007) and filtered with Philosopher (sequential, picked-protein, 1% protein FDR), and TMT-Integrator extracted reporter-ion intensities from the best PSM per peptide at a PSM probability of at least 0.9 and a precursor purity of at least 0.5 with a virtual reference channel. The phosphoproteome search differed in a precursor tolerance of 10 ppm, a fragment tolerance calibrated to 7 ppm, TMTpro fixed on peptide N-termini, methionine oxidation up to two, phosphorylation of serine, threonine and tyrosine up to three per peptide and at most four variable modifications per peptide. Sites were localized with PTMProphet (Shteynberg et al. 2019; expectation maximization, b ions, minimum probability 0.5), the Philosopher filter was sequential at 1% protein FDR, and TMT-Integrator kept sites with a localization probability of at least 0.75, a PSM probability of at least 0.5 and a precursor purity of at least 0.5. The ubiquitinome search used a precursor tolerance of 10 ppm, a fragment tolerance calibrated to 7 ppm, up to three missed cleavages, TMTpro fixed on peptide N-termini only, and diglycine on lysine, TMTpro on lysine, methionine oxidation and protein N-terminal acetylation as variable modifications with at most four per peptide, with PTMProphet localization and the same TMT-Integrator thresholds. The acetyl-lysine and homocitrulline searches used the ubiquitinome settings with acetyl-lysine and carbamyl-lysine in place of diglycine, at most five variable modifications per peptide and a fragment tolerance calibrated to 5 ppm, and TMT-Integrator was run once per tag.

### 8. Protein adjustment of PTM changes and κ calibration

For site s on protein p the adjusted change was $\Delta\_s = \log_2 FC\_s - \kappa \log_2 FC\_p$, with variance $V\_s = SE\_s^2 + \kappa^2 SE\_p^2$ and effective degrees of freedom by the Welch–Satterthwaite approximation, $\nu = V\_s^2/(SE\_s^4/\nu\_s + \kappa^4 SE\_p^4/\nu\_p)$. The statistic $\Delta\_s/\sqrt{V\_s}$ was evaluated against a t distribution, and p-values were adjusted by Benjamini–Hochberg within each layer's reportable site set. This propagation treats site and protein estimation errors as independent and κ as fixed, and the sensitivity of each layer to κ was assessed as described below. Because the channel-median normalization places site ratios on a relative scale, any shift common to all sites of a layer was assessed separately from raw reporter signal.

Phosphorylation was adjusted with $\kappa = 0.46$. The split-half out-of-sample estimate was 0.465 (95% confidence interval 0.440 to 0.480), passenger-peptide regression gave 0.469 against global-run passengers and 0.503 against the protein model, and gene- and pathway-level estimates were 0.462 and 0.511, estimates that assess complementary aspects of the calibration and share some input measurements. The mean held-out residual was −0.003 $\log_2$ units for the adopted calibration, with a constant κ used throughout this data set. Sites were called changed when their BH-adjusted p was below 0.05 at this coefficient, and 10,566 of the 11,079 called sites kept their sign and BH-adjusted significance at both ends of the range spanned by the estimates, 0.44 and 0.51.

The K-ε-GG coefficient was 0.736, based on three estimates that do not share an assumption, a disjoint peptide-half instrumental-variable fit (0.682 over 40 splits), a held-out-bottle residual-correlation criterion (0.763) and a held-out-bottle instrumental-variable fit (0.763). Quantified sites were called changed when their BH-adjusted p was below 0.05 at this coefficient, as for the phosphosites, and 465 of the 483 called sites kept their sign and BH-adjusted significance at both ends of the estimators' range, 0.682 and 0.763.

Acetyl-lysine changes used $\kappa = 0.727$, the mean of three instrumental-variable estimates ranging from 0.68 to 0.79 in which the protein measurement being corrected is never its own regressor, the single-bottle protein ratio of the proteome experiment as the instrument for the site and protein models, the same with site and protein re-estimated from the other two bottles, and the protein ratio from a random half of each protein's peptides over 40 splits. The site model used the 18 NaCl-control halves of the cyanate experiment across the six culture groups (median residual degrees of freedom 9.2) and the protein model used the global proteome, and 109 of the 113 significant sites retained their sign and significance across κ from 0.18 to 1.

Phosphosite annotations were transferred from EPSD (Lin et al. 2021), PhosPhAt (Durek et al. 2010) and UniProtKB/Swiss-Prot onto the *P. celeri* sites by ortholog alignment (Supplementary Data Set 4).

**9. Ubiquitinome enrichment, censoring test and chain linkages**

K-ε-GG peptides were enriched from 430 µg of each digest in 250 µL of PTMScan HS IAP bind buffer containing 0.01% CHAPS with 5 µL of anti-K-ε-GG magnetic beads (Cell Signaling Technology, Danvers, MA, USA; #59322) following Udeshi et al. (2013), incubated for 1 h at 4 °C and processed on a KingFisher Flex. Beads were washed with 50% acetonitrile in IAP wash buffer and then with PBS containing 0.01% CHAPS. Bound peptides were labeled with 400 µg TMTpro reagent in 100 mM HEPES for 20 min, labeling was quenched with 2% hydroxylamine, and after washing the labeled samples were pooled, eluted twice with 0.15% trifluoroacetic acid for 10 min, cleaned on C18 StageTips and reconstituted in 20 µL of 3% acetonitrile with 0.1% formic acid and 0.01% DDM. The pooled plex was analyzed unfractionated in two acquisitions.

Sites detected in at least two of three samples in one condition and none in the other were assessed by a censoring test run on the un-imputed feature and protein values. A global-proteome peptide covering the modified lysine had to be detected in at least two samples of the condition where the site was absent, which establishes evidence for the protein at the relevant region. For each candidate, the mean $\log_2$ intensity of its strongest feature in the detected condition was returned to the raw reporter scale by removing the channel-normalization offsets, and the expected intensity in the absent condition was that value plus κ times the protein change from the detected to the absent condition, taken from an unmodified covering peptide when available and otherwise from the MSstatsPTM protein estimate. The detection floor was the lowest zero threshold among the three absent-condition channels, the lowest nonzero wild-type reporters lying at approximately 2^9.1 to 2^9.2. A change was classified as exceeding protein-dependent censoring when the expected intensity exceeded the floor by more than $1.96 \times \sqrt{[\mathrm{var(feature)}/n + (\kappa\ \mathrm{SE\_protein})^2]}$, as explained by protein change when the expected intensity was below the floor, and as indeterminate in between, with a floor raised by 0.3 $\log_2$ units and the full protein change evaluated as sensitivities. Complete calls and margins are provided in Supplementary Data Set 4, and in site-display panels complete disappearance required detection in all three high-$CO_2$ samples and none in air, whereas incomplete detection was displayed separately.

Chain linkages were assigned from K-ε-GG PSMs on ubiquitin residues K6, K11, K27, K29, K33, K48 and K63, counting only the ubiquitin moieties of polyubiquitin and ubiquitin fusion proteins and excluding remnant events on NEDD8 or ribosomal tails. Reporter intensities were summed per linkage and channel before estimating the air versus high-$CO_2$ change from the three wild-type vessels, with the whole-ubiquitinome shift of −0.648 $log_2$, obtained by pooling 13,728 K-ε-GG PSMs per channel, as the reference. PSM-based 95% intervals describe measurement precision, whereas vessel-based 95% intervals used a t distribution with two degrees of freedom and describe between-vessel reproducibility, PSMs were not counted as independent biological replicates, and linkages supported by fewer than ten PSMs were marked as low-evidence estimates. Set tests on the quantified arm used fry on the protein-adjusted per-channel abundances averaged per gene with the six-group design, and the binary arm used one-sided Fisher's exact tests against proteins carrying a measured K-ε-GG site, with BH within each test collection, the binary-arm enrichment being descriptive because proteins with more measured sites have more opportunities to contribute a lost site.

#### 10. Acetyl-lysine and homocitrulline analyses

Cell pellets were lysed by bead beating (0.5 mm glass beads, Bullet Blender, Next Advance) in 800 µL of 50 mM HEPES pH 7.2 with protease inhibitors (cOmplete Mini EDTA-free, Roche, Basel, Switzerland) and deacetylase inhibitors (10 mM sodium butyrate, 2 µM SAHA, 10 mM nicotinamide), cleared at 14,000 g for 20 min at 4 °C and quantified by BCA, and the cyanate reactions were sealed with gas-impermeable film. Precipitated proteins were redissolved in 5% SDS, 50 mM triethylammonium bicarbonate with the same deacetylase inhibitors before reduction, alkylation and S-Trap digestion. Acetyl-lysine peptides were enriched from each of the four pooled fractions per plex on a KingFisher Flex with 5 µL of antibody beads (PTMScan HS Acetyl-Lysine Motif kit, Cell Signaling Technology, #46784) in IAP bind buffer with 0.01% CHAPS for 3 h at 4 °C, followed by four washes, elution in 0.15% trifluoroacetic acid and C18 StageTip clean-up. Spectra were searched with a precursor tolerance of 10 ppm and an initial fragment tolerance of 20 ppm calibrated to 5 ppm, allowing up to three missed cleavages, and TMT-Integrator produced site-level reporter-ion tables separately for the acetyl and the homocitrulline tag at PSM probability ≥ 0.5, precursor purity ≥ 0.5 and site localization probability ≥ 0.75, with the best PSM per site, outlier removal and MS1-intensity weighting, with and without per-channel median normalization. Gene-set tests of the acetylome used limma fry (Wu et al. 2010) on the per-channel adjusted site abundances averaged per gene, with the six-group design and the wild-type contrast, on the curated pathway sets of this study (at least five genes), the algal-filtered KEGG pathways and the GO terms, reported as $-\log_{10}$ FDR signed by the fry direction.

In the bicarbonate competition, the two loading references available within the immunoprecipitate, the acetyl-lysine peptides and the co-purified unmodified peptides, disagreed in sign about any shift common to all homocitrulline peptides (median per-lysate difference +0.17 and −0.09 $\log_2$), so no global protection was estimated and each site's ratio was centered on the median of all homocitrulline sites of the same lysate. An additional effect-size subset of the limma-protected sites exceeded the tested-site mean by two standard deviations (0.520 $\log_2$). This descriptive cutoff was reported separately from the primary FDR-based selection. For histone H2B K4, isocitrate dehydrogenase K125 and uroporphyrinogen decarboxylase K279, the ratio of the homocitrulline-bearing peptide form was compared with that of the acetylated form of the same peptide across lysates by paired t-test, on PSM-level intensities normalized per channel to the non-homocitrulline PSMs. Set tests used fry with plex as a covariate on the 829 sites complete across the 18 lysates, with sets mapped through the site's gene. The share of ribosomal proteins, the protein's abundance in the global proteome and its lysine content were compared between classes, and acidic (D, E) and basic (K, R) residues were counted within three positions of each lysine in the complete search proteome (1,361 sites), with Mann–Whitney tests between classes and Spearman correlation against the centered ratio.

For homocitrulline in the unenriched proteome, the global, phosphopeptide-enriched and K-ε-GG-enriched data were re-searched in FragPipe 24.0 with TMTpro on lysine as a variable modification (up to three per peptide), carbamyl (+43.0058 Da) on lysine (up to two) and on protein N-termini, and up to five variable modifications per peptide, with PSMs validated by Percolator 3.7.1 (Käll et al. 2007) and filtered to 1% picked protein-level FDR. Rates are homocitrulline matches per total matches of each layer, and sites were pooled across layers by protein position. Each site quantified in all six wild-type channels (35 sites on 25 proteins) was also tested on its own, as the $\log_2$ ratio of its summed reporter intensity, corrected for channel loading, between the three air and the three high-$CO_2$ vessels by Welch's t-test, either as such or after subtracting the $\log_2$ ratio of the unmodified matches of the same protein. To test whether the homocitrulline signal followed its protein, the site $\log_2$ ratio was regressed on the protein $\log_2$ fold change from the MSstatsTMT proteome model by ordinary least squares, with a t-based 95% confidence interval and t-tests of the slope against 0 and against 1. Because RbcL contributed seven sites, elongation factor 1α four and histone H3.2 two, uncertainty was also assessed by a case bootstrap that resampled the 25 proteins with replacement together with all their sites (10,000 draws, percentile interval). The influence of individual proteins was assessed by omitting each protein in turn, with the fit excluding all RbcL sites shown in Supplementary Figure 2 and the retained fits provided in Supplementary Data Set 4.

For acetylation-site conservation, *P. celeri* and *Chlamydomonas reinhardtii* (CC-4532 v6.1) proteins were paired by reciprocal best hits with DIAMOND (very-sensitive mode, e-value $10^{-5}$; Buchfink et al. 2021) and aligned globally with BLOSUM62 (gap open −11, extension −1; Biopython, Cock et al. 2009), and each acetylated lysine was transferred to its aligned residue. Chlamydomonas acetylation sites (Füßl et al. 2022; 1,378 sites) were relocated into v6.1 by their 31-residue sequence windows (1,261 placed). Conservation of a site was tested against every other conserved lysine of the same protein pairs by the Cochran–Mantel–Haenszel test stratified by protein pair, with a sensitivity analysis restricted to pairs with a unique optimal alignment. RbcL positions follow the spinach numbering (UniProt P00875), to which the *P. celeri* sequence aligns without gaps at 88% identity.

### 11. Metabolomics and lipidomics acquisition

For GC-MS, extracts were derivatized by methoximation and trimethylsilylation (Kim et al. 2015), and 1 µL was injected splitless at 250 °C onto an HP-5MS column (30 m × 0.25 mm × 0.25 µm) on an Agilent 7890A coupled to a 5975C detector, with helium at 0.71 mL $min^{-1}$, the transfer line at 280 °C, the source at 250 °C and the quadrupole at 150 °C. The oven was held at 60 °C for 1 min, raised at 10 °C $min^{-1}$ to 325 °C and held for 10 min, and spectra were scanned over m/z 50 to 600 after a 6-min solvent delay. C8 to C28 fatty acid methyl esters (Sigma-Aldrich, St. Louis, MO, USA) were analyzed in parallel for retention-index calibration. Metabolite Detector (Hiller et al. 2009) performed deconvolution, retention-index calculation and alignment, spectra were matched against the PNNL-augmented Agilent metabolomics library (Kind et al. 2009) with unmatched compounds searched against NIST20 and Wiley11, and identification and quantification ions were manually validated.

For HILIC LC-MS/MS, 10 µL was injected onto an ACQUITY UPLC BEH HILIC column (2.1 × 100 mm, 1.7 µm, Waters) at 50 °C on a Vanquish Flex UHPLC. Solvent A was 10 mM ammonium acetate and 0.05% ammonium hydroxide in 95:5 water:acetonitrile, and solvent B was 0.05% ammonium hydroxide in acetonitrile, B being the complement of A in the following program.

| Time (min) | A (%) | Flow (µL $min^{-1}$) | Segment |
|---|---|---|---|
| 0–6.0 | 5 → 63 | 300 | Gradient |
| 6.0–7.0 | 63 | 300 | Hold |
| 7.0–7.1 | 63 → 5 | 300 | Return |
| 7.1–7.2 | 5 | 300 → 500 | Increase flow |
| 7.2–9.5 | 5 | 500 | Flush |
| 9.5–9.7 | 5 | 500 → 300 | Decrease flow |
| 9.7–12.0 | 5 | 300 | Re-equilibrate |

The Orbitrap Eclipse heated electrospray source operated in negative mode at 3.0 kV, with the capillary at 350 °C, an S-lens RF level of 30 and the auxiliary-gas heater at 250 °C. Full scans covered m/z 67 to 1000 at 120,000 resolving power (FWHM at m/z 200), and data-dependent MS2 used standard AGC, a 100-ms maximum injection time, 1.6 m/z isolation and a loop count of 12, with HCD spectra at 15,000 resolving power and stepped normalized collision energies of 20, 30 and 40 and CID spectra in the ion trap at 30% collision energy and 10-ms activation time. Compound Discoverer 3.3 used adaptive-curve alignment with a maximum retention-time shift of 0.5 min, a mass tolerance of 5 ppm and a minimum peak intensity of $2.5 \times 10^5$, and annotations used isotopic patterns, retention time and MS1/MS2 evidence and were manually reviewed. The four EMSL annotation categories defined in the main Methods are retained as facility metadata rather than asserted to be equivalent to MSI identification levels, and the reported HILIC results are from negative mode.

Metabolite PQN corrected a sample-intensity spread of approximately three $\log_2$ units, with 40 of 99 metabolites passing BH-adjusted $p < 0.05$ after PQN compared with two without normalization. Restricting the model to wild-type samples gave 38 significant metabolites, while the lipidome yielded 104 significant species without normalization and 121 with PQN (Supplementary Data Set 2).

Dried organic MPLEx extracts were resuspended in 200 µL of 10% chloroform in methanol (v/v), and 10 µL was loaded onto a guard column (ACQUITY UPLC CSH C18 VanGuard pre-column, 130 Å, 1.7 µm, 2.1 mm × 5 mm, Waters) ahead of a reversed-phase column (ACQUITY UPLC CSH C18, 130 Å, 1.7 µm, 3.0 mm × 150 mm, Waters, facility column identifier WCSH415413) on an ACQUITY UPLC H-Class system (Waters) held at 50 °C. Mobile phase A was acetonitrile:water (40:60, v/v) and mobile phase B was acetonitrile:isopropanol (10:90, v/v), each with 10 mM ammonium acetate, delivered at 300 µL $min^{-1}$ over the 21-min gradient below within a 25-min acquisition.

| Time (min) | B (%) | Flow (µL $min^{-1}$) | Segment |
|---|---|---|---|
| 0–1.0 | 40 → 62 | 300 | Gradient |
| 1.0–4.0 | 62 → 66 | 300 | Gradient |
| 4.0–9.0 | 66 → 78 | 300 | Gradient |
| 9.0–11.0 | 78 → 87 | 300 | Gradient |
| 11.0–15.0 | 87 → 99 | 300 | Gradient |
| 15.0–21.0 | 99 | 300 | Hold |

The Orbitrap Fusion Lumos (Thermo Fisher Scientific) heated electrospray source operated at 3,500 V in positive mode and 3,400 V in negative mode, with sheath, auxiliary and sweep gas at 50, 10 and 1 arbitrary units, an ion-transfer-tube temperature of 300 °C and a vaporizer temperature of 350 °C. MS1 scans covered m/z 200 to 1800 at 120,000 resolution with an automatic gain control (AGC) target of $4 \times 10^5$ and a 50-ms maximum injection time. Precursors above $2.5 \times 10^4$ counts were selected within a 1-s cycle time and excluded for 4 s after two selections. Each selected precursor was fragmented with 1 m/z isolation by HCD at stepped normalized collision energies of 25, 30 and 35 (Orbitrap detection at 7,500 resolution, AGC target $1 \times 10^5$, 50-ms maximum injection time) and by CID at 38% collision energy for 10 ms (ion-trap detection, AGC target $2 \times 10^4$, 35-ms maximum injection time). A pooled quality-control sample was injected four times in each mode alongside three process blanks. Lipids were identified with LIQUID (Lipid Informed Quantitation and Identification; Kyle et al. 2017) by examining the diagnostic ion fragments and acyl-chain fragments in each tandem spectrum together with the mass error of the precursor, its isotopic profile and its extracted-ion chromatogram. A target database of the identified lipid names, retention times and observed m/z was then used to identify features across all runs, with MZmine 2 (Pluskal et al. 2010) aligning and gap-filling the individual runs by matching unidentified features to their identified counterparts, and all aligned features were manually verified before peak intensities were exported. No internal standards were included, and the exported values are relative peak intensities that support fold-change comparisons within a lipid class but not comparisons of amount between classes, whose ionization efficiencies differ. The facility inventory of 307 lipid names in 16 subclasses contained 285 measured rows, of which 14 positive-mode pairs of positional isomers (same class and sum composition, same retention time, identical intensities in every wild-type sample, QC pool and blank) shared one chromatographic peak and were counted once under a joined name, the convention the inventory itself uses for 26 other unresolved co-eluting pairs, leaving 271 measured features. Of these, 32 failed the 30% relative-standard-deviation threshold, and the final wild-type detection and QC filters retained 237 species.

#### 12. RNA-seq processing pipeline

Annotation and genome. The two haplotype GFF3 exports were merged after feature sorting, URL decoding, tRNA identifier de-duplication and exon generation into one unified GFF3 of 13,833 genes (13,746 nuclear, 56 chloroplast, 31 mitochondrial) on 32 scaffolds, with one transcript per gene and Parent as the transcript tag. This annotation, unified.gff3, was used for the STAR index, whereas fragment counting used the revised unified_v3.gff3 annotation (file checksums in Supplementary Data Set 2). Allelic pairs between the haplotypes were defined by whole-chromosome nucmer alignment (MUMmer4; Marçais et al. 2018), giving 6,351 pairs at 98.7% high confidence and 1,044 unpaired genes (639 in haplotype p0, 405 in p1).

Trimming. fastp 1.0.1 (Chen et al. 2018) with paired-end adapter detection, overlap-based base correction, sliding-window trimming from both read ends (window 4, mean quality 20), a qualified-base threshold of Phred 20 with at most 30% unqualified bases per read, and a minimum read length of 36 nt. Of 543,173,673 read pairs, 539,052,835 (99.24%) passed.

Index and alignment. The STAR 2.7.11b (Dobin et al. 2013) index was built from the unified genome and GFF3 with sjdbOverhang 149 and genomeSAindexNbases 11. Reads were aligned in two-pass mode (twopassMode Basic) with outFilterMultimapNmax 200 and winAnchorMultimapNmax 200, so that a read matching both haplotypes is retained, and outSAMmultNmax 1, so that one alignment per read is written, with NH, HI, AS, NM and MD attributes and coordinate-sorted BAM output. Alignment rates were about 40% unique and 53% multi-mapped, the multi-mapped fraction being the allelic reads.

Alignment quality control. RSeQC 5.0.3 (Wang et al. 2012) modules bam_stat, infer_experiment, inner_distance, junction_saturation, read_distribution, read_duplication, geneBody_coverage and tin were run on every BAM, and reports were collected with MultiQC. infer_experiment on 200,000 sampled pairs per library assigned 97.82% of pairs to the reverse-stranded orientation, which set the strand option of the counting step.

Counting and allelic collapse. featureCounts 2.1.1 (Liao et al. 2014) counted fragments per transcript (exon features grouped by Parent) in paired-end mode with reverse strandedness (-s 2) and multi-mapping fragments counted once (-M). Because each read carries one alignment, an allelic read is counted at one of the two haplotypes and the pair sum is complete after collapse. Transcript counts were mapped to gene identifiers, the two alleles of each pair were summed into one gene, unpaired genes were kept as single genes, and chloroplast and mitochondrial genes were removed, giving the gene-level matrix.

Differential expression. Genes with fewer than 10 total counts across the 18 libraries were removed, leaving 7,352. DESeq2 1.42.0 (Love et al. 2014) was fitted with the design ~ group, one coefficient per genotype-and-condition group, and the contrast wild type in air against wild type at high $CO_2$ was tested at alpha 0.05 with the default independent filtering and Benjamini–Hochberg adjustment. Fold changes were shrunk with apeglm (Zhu et al. 2019) on the corresponding coefficient, and the unshrunken Wald statistic and adjusted p-value were kept alongside the shrunken $\log_2$ fold change. Genes with adjusted p below 0.05 and an absolute shrunken $\log_2$ fold change of at least 1 were called differentially expressed.

Enrichment. Over-representation analysis used clusterProfiler enricher (Wu et al. 2021) on GO biological process, GO molecular function and KEGG terms with the 7,352 tested genes as universe, term sizes 5 to 500 and an adjusted p cutoff of 0.05. Gene-set enrichment used fgsea (Korotkevich et al. 2021) on genes ranked by the DESeq2 Wald statistic with term sizes 10 to 500 for GO and 10 to 1,000 for KEGG. Human-disease and non-plant terms were removed before testing. The curated pathway sets were tested with fry and camera (limma; Wu et al. 2010; Wu and Smyth 2012; Ritchie et al. 2015) on $\log_2$ counts per million with the same 18-sample design and contrast.

### Supplementary Figure 1

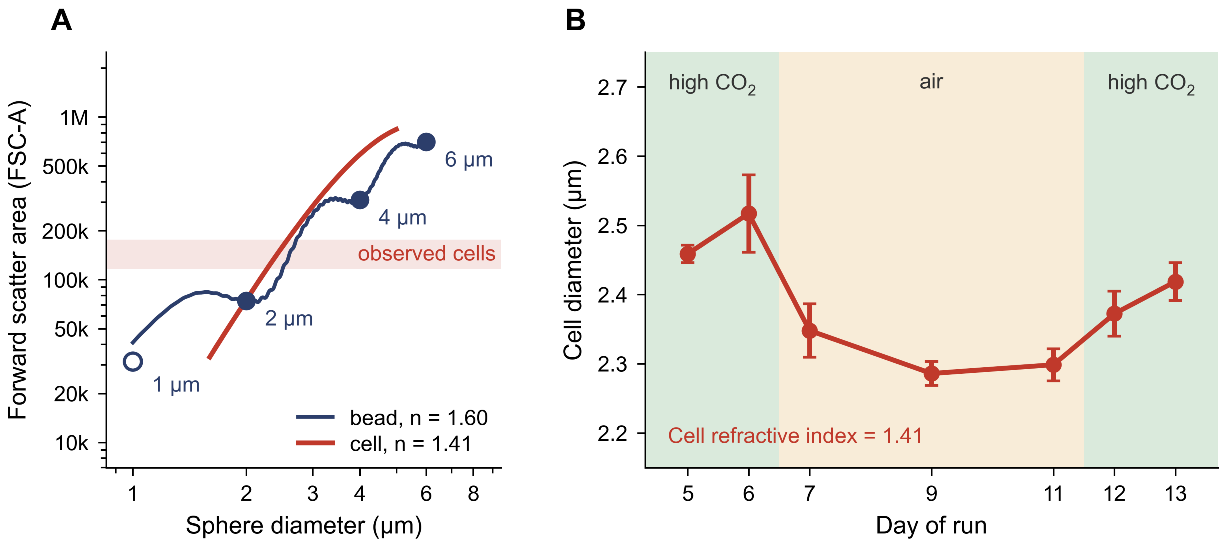


Supplementary Figure 1. Bead calibration and Mie inversion of cell diameter from flow cytometry. A) Forward scatter area (FSC-A) against sphere diameter on logarithmic axes. The dark line is the Mie-theory curve for polystyrene (refractive index 1.60) under the collection window fitted to the 2, 4 and 6 µm beads (filled circles, modes located by kernel density estimation), whereas the 1 µm bead (open circle) sits at the acquisition threshold and was not used in the fit. The red curve is the Mie curve for a homogeneous cell at a refractive index of 1.41 under the same window with the chlorophyll absorption term set to 0.003, and the shaded band spans the median FSC-A of the 27 retained cell samples. The polystyrene curve oscillates because a sphere that refracts light much more strongly than water produces Mie resonances in forward scatter at these sizes, whereas the cell curve is smoother because the refractive index of the cell is close to that of water. B) Cell diameter by day of the second run under high $CO_2$, air and restored high $CO_2$ (shading). Red circles and error bars show the mean and standard deviation over four vessels, with three vessels on Day 9. Diameters were calculated at a cell refractive index of 1.41. Values are given in Supplementary Data Set 1 and support the cell-size measurements in Figure 1.

### Supplementary Figure 2

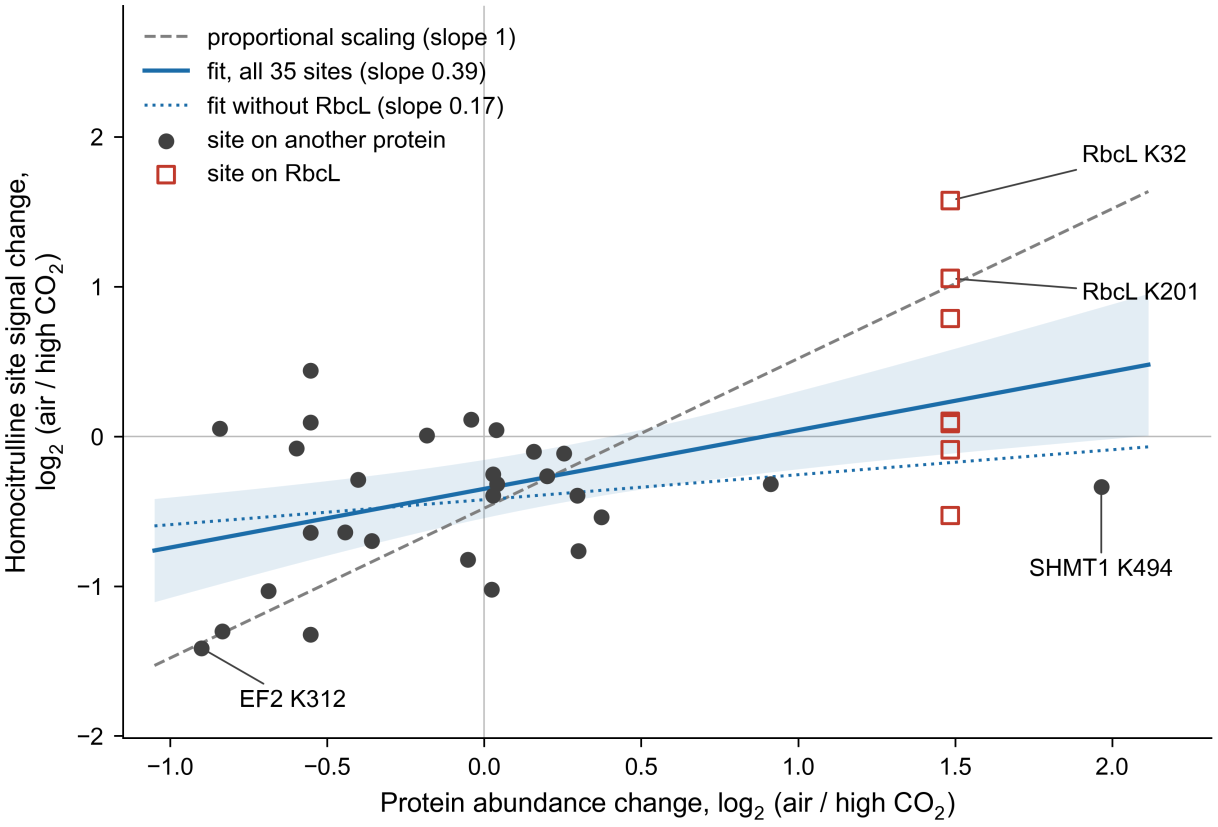


Supplementary Figure 2. Homocitrulline site signal changes less than the abundance of the protein that carries it. Supporting the homocitrulline-persistence analysis in the Results, the $\log_2$ change in air relative to high $CO_2$ of the reporter-ion signal of each homocitrulline site is plotted against the $\log_2$ change of its protein from the MSstatsTMT proteome model, for the 35 sites on 25 proteins that were quantified in all six wild-type channels of the unenriched proteome re-searches. The seven RbcL sites are open squares, all other sites are filled circles, and four sites are labeled. The dashed gray line is proportional scaling (slope 1). The solid blue line is the ordinary least-squares fit through all 35 sites (slope 0.39, 95% confidence interval 0.16 to 0.62, t-test with 33 degrees of freedom, $p = 0.0014$ against a slope of 0 and $p = 5 \times 10^{-6}$ against a slope of 1) with its pointwise 95% confidence band, whereas the dotted line is the fit without the RbcL sites (slope 0.17, 95% confidence interval −0.15 to 0.48). Site values, protein changes and every sensitivity analysis of the slope are given in Supplementary Data Set 4.
